# Gut microbiome dysbiosis is associated with elevated toxic bile acids in Parkinson’s disease

**DOI:** 10.1101/2020.09.26.279851

**Authors:** Peipei Li, Bryan A. Killinger, Ian Beddows, Elizabeth Ensink, Ali Yilmaz, Noah Lubben, Jared Lamp, Meghan Schilthuis, Irving E. Vega, Markus Britschgi, J. Andrew Pospisilik, Patrik Brundin, Lena Brundin, Stewart Graham, Viviane Labrie

## Abstract

The gut microbiome can impact brain health and is altered in Parkinson’s disease (PD) patients. Here, we investigate changes in the functional microbiome in the appendix of PD patients relative to controls by metatranscriptomic analysis. We find microbial dysbiosis affecting lipid metabolism, particularly an upregulation of bacteria responsible for secondary bile acid synthesis. Proteomic and transcript analysis corroborates a disruption in cholesterol homeostasis and lipid catabolism. Bile acid analysis reveals an increase in microbially-derived, toxic secondary bile acids. Synucleinopathy in mice induces similar microbiome alterations to those of PD patients. The mouse model of synucleinopathy has elevated DCA and LCA. An analysis of blood markers shows evidence of biliary abnormalities early in PD, including elevated alkaline phosphatase and bilirubin. Increased bilirubin levels are also evident before PD diagnosis. In sum, microbially-derived toxic bile acids are heightened in PD and biliary changes may even precede the onset of overt motor symptoms.

## Introduction

Parkinson’s disease (PD) is a common neurodegenerative disease that is clinically characterized by motor and non-motor symptoms. Some non-motor features of PD begin many years before the onset of motor symptoms, during the prodromal phase of PD (Kalia and Lang, 2015). One of the first prodromal symptoms is constipation, pointing toward an early involvement of the gastrointestinal (GI) tract (Kalia and Lang, 2015). Aggregated α-synuclein α-syn), a pathological hallmark of PD, is apparent in the GI tract of prodromal PD patients (Hilton et al., 2014; Stokholm et al., 2016). α-Syn aggregates in the gut of experimental models have been reported to propagate to the brain and induce nigral neurodegeneration and PD-like motor and non-motor dysfunctions (Kim et al., 2019; Van Den Berge et al., 2019). Recently, the vermiform appendix has been implicated as one GI tract location that could contribute to the initiation of PD pathogenic processes (Killinger et al., 2018). The appendix contains an abundance of aggregated α-syn, particularly in enteric nerves, with PD patients having substantially higher amounts of these aggregates (Killinger et al., 2018; Stokholm et al., 2016). Removal of the appendix was associated with a decreased risk for PD in some, but not all, epidemiological studies (Killinger et al., 2018; Liu et al., 2020; Marras et al., 2016; Mendes et al., 2015; Svensson et al., 2016). This signifies that the appendix may be an important tissue to study to advance our understanding of some of the earliest events in PD pathogenesis.

The appendix is an immunological organ that also acts as a storehouse for the gut microbiome (Donaldson et al., 2016; Killinger and Labrie, 2019). The gut microbiome and its metabolites are increasingly being recognized as crucial for brain health (Fung et al., 2017). Numerous studies support that there are microbiome changes in the stool of PD patients (Bedarf et al., 2017; Heinzel et al., 2020; Perez-Pardo et al., 2019; Scheperjans et al., 2015). The appendix contains a rich microbial biofilm, which differs from that of the rectum and stool (Jackson et al., 2014; Tytgat et al., 2019). Importantly, the appendix has an anatomically shielded microbiome and is capable of modulating the microbiome in the rest of the large intestine (Masahata et al., 2014; Sanchez-Alcoholado et al., 2020). Consequently, changes in the appendix microbiome may have a widespread effect on the microbial flora of the intestine. Further, inflammation in the periphery and the brain has been proposed to have a central role in PD, and the microbiome and the host immune system has a bidirectional effectual relationship (Fung et al., 2017; Johnson et al., 2019). In the appendix, microbial metabolites have direct access to the immune cells within the lymphoid follicles and may thereby modify inflammatory responses and immunity (Bachem et al., 2019; Chun et al., 2019; Schulthess et al., 2019). Microbial metabolites can also alter brain inflammation and motor impairments in PD models (van Kessel and El Aidy, 2019). Thus, dysregulation of the appendix microbiome may be involved in PD, but this has yet to be examined.

One important function of the gut microbiome is its involvement in the biotransformation of bile acids. Bile acids are amphipathic molecules that aid in the absorption of dietary lipids and also affect glucose homeostasis, inflammation, gastrointestinal functions, as well as blood–brain– barrier integrity and signaling in the brain (Hegyi et al., 2018; McMillin and DeMorrow, 2016; Wahlström et al., 2016). In the liver, primary bile acids — cholic acid (CA) and chenodeoxycholic acid (CDCA) — are synthesized from cholesterol. Bile acids are stored and concentrated in the gallbladder before their release into the small intestine. Most primary bile acids are reabsorbed in the distal ileum for transport back to the liver as a part of the enterohepatic circulation (Wahlström et al., 2016). The remaining primary bile acids that enter the large intestine are converted by the microbiome into secondary bile acids — deoxycholic acid (DCA), lithocholic acid (LCA), and to a lesser extent, ursodeoxycholic acid (UDCA). Secondary bile acids are produced solely by bacteria, largely by those in *Clostridium* clusters *XIVa* and *XI* of the *Firmicutes* phylum (Wahlström et al., 2016). LCA and DCA are hydrophobic bile acids that are cytotoxic at elevated physiological concentrations (Chen et al., 2019a; Pavlidis et al., 2015). LCA, the most toxic bile acid, is highly insoluble, which limits its reabsorption, and it is primarily excreted in stool. For DCA, the second most cytotoxic bile acid, approximately 50% is passively reabsorbed in the large intestine, and it returns via the enterohepatic circulation for detoxification in the liver. Increases of LCA and DCA have been implicated in intestinal inflammation, liver injury, cholestasis, and gallstone formation (Berr et al., 1994; Chen et al., 2019a; Hang et al., 2019; Song et al., 2020). Whether there are hydrophobic bile acid changes or bile-related illnesses that impact PD risk is unknown.

Here, we performed a metatranscriptomic analysis of the appendix microbiome in PD patients and controls. In parallel, we performed an in-depth analysis of the appendix microbiome of a mouse model of synucleinopathy and examined the effects of gut inflammation. The microbial taxa and pathway alterations we identified led us to an analysis of bile acids in PD patients and in the mouse model, where we identified increases in the hydrophobic bile acids LCA and DCA. This was consistent with proteomic and transcript analyses revealing perturbations in lipid metabolism and uptake in the PD gut. In an analysis of liver and gallbladder disease blood markers, we found evidence of biliary abnormalities in PD, including elevated bilirubin, which was associated with a worsening of motor dysfunction. In at-risk individuals, increased bilirubin levels preceded PD diagnosis, indicating that bile-related abnormalities may represent upstream events in PD pathogenesis. Thus, microbiome-mediated changes in hydrophobic bile acids and a disruption of biliary function may be involved in the development of PD.

## Results

### Microbiome changes in the PD appendix and in response to synucleinopathy and gut inflammation

We performed a comprehensive analysis of the microbiome in appendix tissue of PD patients and controls using metatranscriptomic sequencing. Metatranscriptomic analysis profiles the functionally active microbiome. Our analysis examined the appendix microbiome of 12 PD patients and 16 controls, and adjusted for age, sex, postmortem interval, and RIN (tissue quality). Postmortem appendix tissue was from PD patients with confirmed brain Lewy pathology (PD Braak stage ≥ 3), while controls had no brain Lewy pathology (Supplementary File 1). We had on average 14,288,947 ± 5,008,507 reads per sample for microbiome analysis after data quality control. Microbial taxa were identified using MetaPhlAn2, and we found transcripts for 65 genera, 37 families, 20 orders, 15 classes, and 9 phyla in the human appendix. Overall, we did not find changes in the richness of microbiome communities between PD patients and controls at any taxonomic level (alpha diversity, Shannon index; Figure S1). The appendix of PD patients and controls also had a similar overall microbial community composition (beta-diversity, Whittaker index and NMDS ordination; Figure S1). The most abundant bacteria in the appendix of PD patients and controls were *Lachnospiraceae*, *Ruminococcaceae*, *Porphyromonadaceae*, *Enterobacteriaceae*, and *Bacteroidaceae*, together accounting for 68.6% of the relative family abundance (Figure S1). As a whole, the appendix microbial community identified in our metatranscriptomic analysis was similar to that found in surgically-isolated, healthy appendix tissues (Jackson et al., 2014) (order level: R=0.91, *p*<10^-14^; family level: R=0.26, *p*<0.05; Pearson’s correlation).

In an analysis examining the abundance of microbial taxa, differences between the appendix microbiome of PD patients and that of controls were observed at all taxonomic levels (FDR *q*<0.05, analysis using metagenomeSeq (zero-inflated Gaussian mixture model) and adjusting for age, sex, postmortem interval and RIN; **Figure 1A**; Supplementary File 2). The most significant change in the PD appendix microbiome relative to controls was in the order of *Clostridiales*, particularly an increase in *Peptostreptococcaceae* (*Clostridium* cluster *XI*; *q*<0.05 observed across family and genus) and in *Lachnospiraceae* (*Clostridium* cluster XIVa; *q*<0.05 family). The appendix of PD patients also had a prominent increase in *Burkholderiales* (family *Burkholderiaceae*, genus *Burkholderia*; *q*<0.05 observed across class, order, family, and genus) and a decrease in *Methanobacteriales* (family *Methanobacteriaceae*, genus *Methanobrevibacter*; *q*<0.05 across all taxonomic levels). Furthermore, the PD appendix had decreases of the genera *Odoribacter*, *Clostridium*, *unclassified Sutterellaceae*, and *Escherichia* (*q*<0.05 genus).

**Figure 1:**
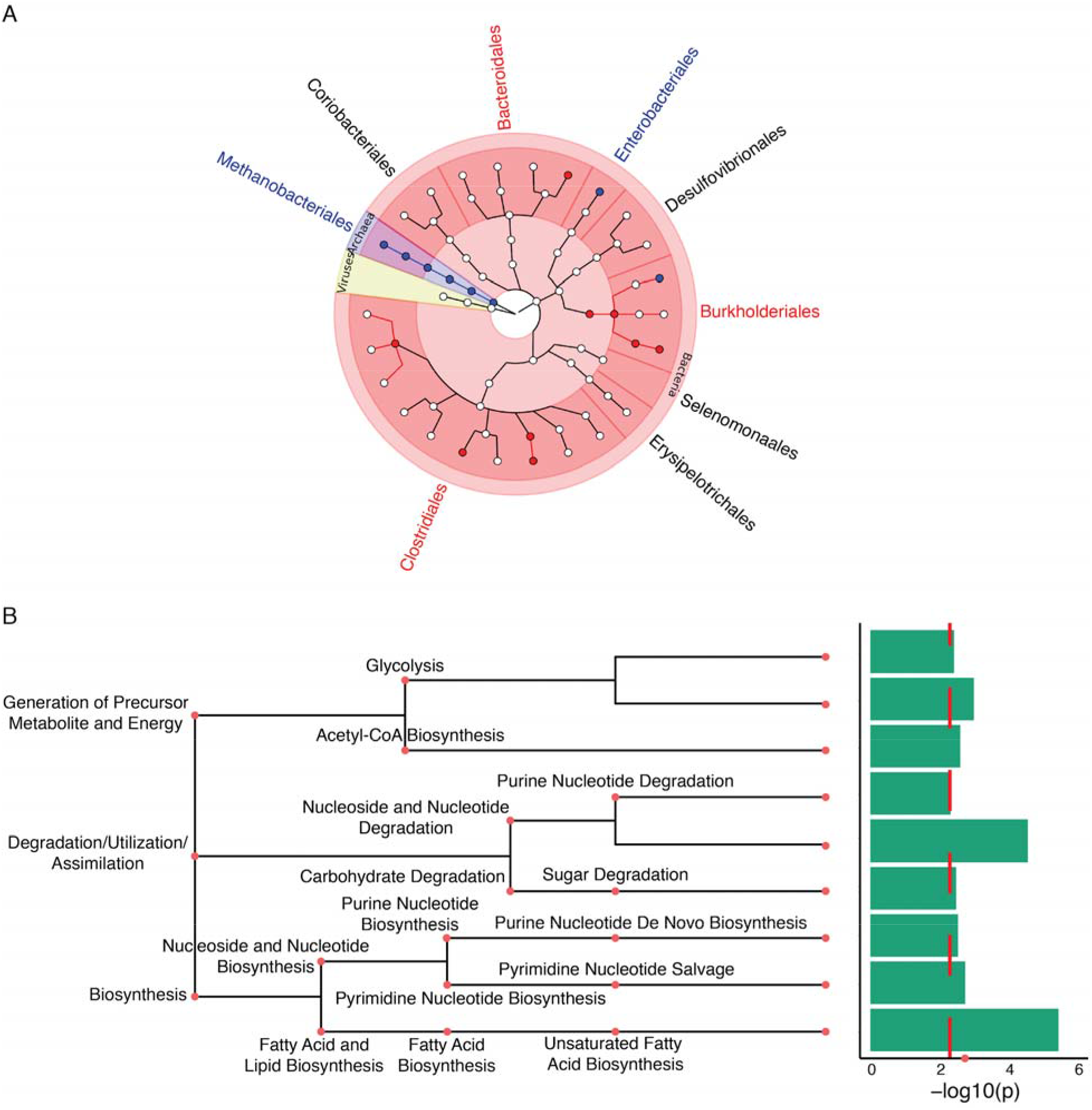
The PD appendix exhibits shifts in the active microbiome that affect lipid metabolism. Metatranscriptomic analysis was used to determine changes in the functional microbiome in the appendix of PD patients (n=12 PD, 16 controls). (**A**) Microbiome changes in the PD appendix. Metatranscriptome data were analyzed by MetaPhlAn2 and a zero-inflated gaussian mixture model in metagenomeSeq, adjusting for age, sex, RIN, and postmortem interval. Results are displayed using GraPhlAn, showing the taxonomic tree with kingdom in the center, and branching outwards to phylum, class, order, family, and genus. Microbial taxa highlighted in red are increased in PD, and blue are decreased in PD (*q*<0.05, metagenomeSeq). (**B**) Microbiome metabolic processes altered in the PD appendix. Top microbial pathways altered in PD identified by HumanN2. Red dashed line denotes *q*<0.1 pathways as determined by metagenomeSeq.

To explore factors contributing to microbiome changes in PD and to control for the environment and diet, we explored appendix microbiome changes in a PD mouse model of synucleinopathy exposed to gut inflammation (**Figure 2A**). In mice, the cecal patch is the appendix equivalent. We examined the cecal patch microbiome in a synucleinopathy model: mice overexpressing human α-syn with the heterozygote A30P mutation) (A30P αsyn mice) (Kahle et al., 2000). Gut inflammation was induced by a chronic treatment of dextran sulfate sodium (DSS), a widely used model of ulcerative colitis (Chassaing et al., 2014). Adult A30P α-syn and wild-type mice were exposed to increasing concentrations of DSS over four weeks (2.5% to 4% DSS), followed by a four-week recovery period. The cecal patch microbiome was profiled by 16S rRNA sequencing in A30P α-syn and wild-type mice previously exposed to DSS or water (n=8-11 mice/group; average 218,198 ± 5752 reads/sample; Supplementary File 3). There were 124 genera, 53 families, 29 orders, 18 classes, and 11 phyla identified. We observed no differences in the richness of the microbial communities between the mouse groups (alpha diversity, Shannon index; Figure S2). The mouse groups also had a similar overall microbial community composition (beta-diversity, NMDS ordination; Figure S2). The microbial community in the mouse cecal patch was similar overall to that of the human appendix in the metatranscriptomic analysis above (order level: R=0.88, *p*<10^-10^; family level: R=0.50, *p*<10^-4^; Pearson’s correlation).

**Figure 2:**
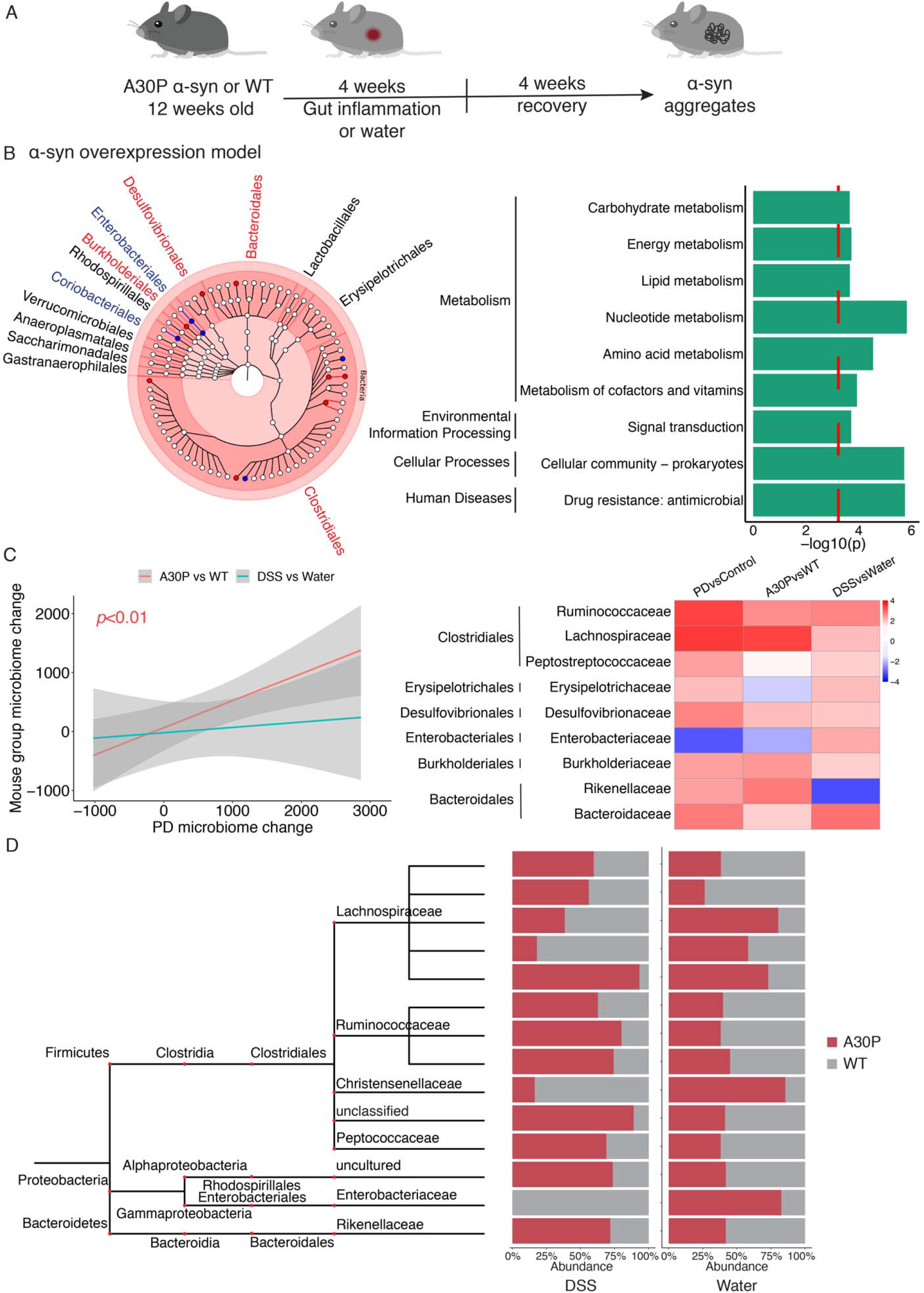
Microbiome changes induced by synucleinopathy and gut inflammation in the mouse cecal patch. The microbiome was profiled by 16S rRNA sequencing in the cecal patch of a mouse model of synucleinopathy, A30P α-syn overexpressing mice. In addition, microbiome changes in responses to gut inflammation by prior DSS colitis exposure was determined in the A30P α-syn and wild-type mice. The 16S rRNA data were analyzed using QIIME2 and a zero-inflated gaussian mixture model in metagenomeSeq, adjusting for covariates (sex and DSS exposure or genotype) was used to identify differences in the microbiome (*q*<0.05, metagenomeSeq). Microbial metabolic pathway changes were determined by PICRUSt2. n=40 mice, 8 A30P/DSS, 10 A30P/water, 11 WT/DSS, 11 WT/water. (**A**) Schematic of experimental design to examine microbiome changes in A30P α-syn or wild-type mice exposed to DSS colitis or water. (**B**) Microbiome changes altered in response to synucleinopathy. Taxonomic tree showing changes in microbial taxa (left panel) and related microbial functional pathways (right panel) in the A30P α-syn overexpression model, relative to wild-type mice. Taxonomic tree built using GraPhlAn, with kingdom in the center, and branching outwards to phylum, class, order, family, and genus. Microbial taxa highlighted in red are in A30P α-syn mice relative to wild-type mice, and blue are decreased in A30P α-syn mice. Top microbial pathways altered by synucleinopathy are shown. Red dashed line denotes *q*<0.1 pathways determined by metagenomeSeq. (**C**) The mouse model of synucleinopathy recapitulates microbiome changes in the PD appendix. Left panel: Correlation of microbiome changes at family level in the PD appendix with those altered in the cecal patch of the mouse model of synucleinopathy and gut inflammation. *p*<0.01 represents a significant correlation between microbiome changes in the PD appendix and those occurring in response to A30P α syn overexpression. Right panel: Heatmap demonstrating microbiome differences at the family level in the PD appendix and in the cecal patch of mice with A30P α-syn overexpression or in response to DSS colitis. Microbial families existing in both humans and mice are shown. Red signifies an increase in the microbial taxa in PD or mouse model, and blue signifies a decrease. (**D**) Microbiota changes in the A30P α-syn mouse cecal patch in response to gut inflammation. Abundance (percentage) of each microbial genera altered in synucleinopathy model by DSS colitis with nominal *p*<0.05. Comparison between the DSS panel (left) and water panel (right) demonstrates changes in microbiota abundances in A30P α-syn mice in response to DSS, relative to wild-type mice. Taxonomic tree shows from phylum to family level.

We investigated differences in the abundance of microbial taxa in the cecal patch that were induced by synucleinopathy. In mice overexpressing A30P α-syn, the most significant changes were within the order *Clostridiales*, including an increase within the *Lachnospiraceae* family (*A2* genus), in the *Christensenellaceae* family, and in unclassified *Clostridiales* (*q*<0.05, metagenomeSeq; **Figure 2B**; Figure S3; Supplementary File 4). However, there were also decreases in *Clostridiales* on the genus level (*q*<0.05 genera within the family *Ruminococcaceae* and *Family XIII*). Similar to the PD appendix, A30P α-syn overexpressing mice had an increase in *Burkholderiales* relative to wild-type mice (family *Burkholderiaceae, q*<0.05; **Figure 2B**; Figure S3; Supplementary File 4). Also, these mice had a decrease in *Enterobacteriales* (family *Enterobacteriaceae*, *q*<0.05 observed in order and family) and *unclassified Bacteroidales* (*q*<0.05 genus), as observed in PD. Moreover, we found that microbial changes at the family level were significantly correlated between the synucleinopathy model and the PD appendix (R=0.8; *p*<0.01, Pearson’s correlation; **Figure 2C**). Overall, synucleinopathy in mice induces microbial changes that significantly mimic those of the PD appendix.

We next examined the effects of gut inflammation on microbial composition in the mouse cecal patch. Prior DSS exposure induced a prominent increase in *Coriobacteriales* (across all taxonomic levels), decreases in *Bacteriodales* (family and genus), an increase within *Desulfovibrionales* (genus), and mixed effects in family and genera within *Clostridiales* (*q*<0.05, metagenomeseq; Figure S4; Supplementary File 5). However, in general, DSS exposure in mice did not result in changes in the microbiome that resembled those of the PD appendix (R=0.19; *p*=0.63, Pearson’s correlation; **Figure 2C**). In the A30P mouse model of synucleinopathy, DSS exposure intensified changes within the order *Clostridiales* (*q*<0.05 family and genus level; Figure S5; Supplementary File 6). The A30P α-syn mice, relative to wild-type mice, had a greater accumulation of genera within *Clostridiales* in response to DSS colitis, including an increase in *Ruminococcaceae* (*q*<0.05 genus; **Figure 2D**). Also, A30P α-syn mice had a strong decrease in *Enterobacteriales* following DSS exposure in comparison to wild-type mice (family *Enterobacteriaceae, q*<0.05 order and family; **Figure 2D**; Figure S5). In sum, synucleinopathy in mice partly recapitulates microbiome changes observed in the PD appendix, and experimentally-induced gut inflammation amplifies some of these microbiome alterations.

Next, we examined whether there were microbial metabolic pathways altered in common between the PD appendix and the mouse model of synucleinopathy and gut inflammation. In the PD appendix, the most significant microbial pathway change was a loss of fatty acid metabolism (*q*<0.1, HUMANn2; **Figure 1B**; Supplementary File 2). Likewise, there was a decrease in lipid metabolism in the A30P α-syn overexpressing mice when compared to wild-type mice (*q*<0.1, PICRUSt2 and metagenomeSeq; **Figure 2B**; Supplementary File 4). Synucleinopathy combined with DSS inflammation also decreased lipid metabolism pathways (*q*<0.1, Figure S5; Supplementary File 6). Microbial shifts resulting from synucleinopathy and gut inflammation in mice also affected other metabolic processes, including carbohydrate, nucleotide, amino acid, and vitamin metabolism, and altered antimicrobial drug resistance (*q*<0.1, PICRUSt2 and metagenomeSeq; **Figure 2B**; Figure S4, S5). Thus, dysregulation of lipid metabolism is a microbial community pathway alteration that is shared between the PD appendix and a mouse model of synucleinopathy and gut inflammation.

Considering the prominent disruption of lipid metabolism pathways in the microbiome of PD patients, we investigated whether there were corresponding changes in the host. We analyzed the human proteome in appendix tissue of PD patients and controls. Pathway analysis of the PD appendix proteome showed a reduction in proteins affecting lipid metabolism (*q*<0.05 pathways, hypergeometric test; **Figure 3**; Supplementary File 7). There was also a dysregulation of pathways involved in protein localization, antigen presentation, glycolysis, and immune activity in the PD appendix (*q*<0.05 pathways, hypergeometric test; **Figure 3**). Overall, human proteomic changes affecting lipid homeostasis in the PD appendix corroborate the microbial pathway alterations affecting lipids in patients.

**Figure 3:**
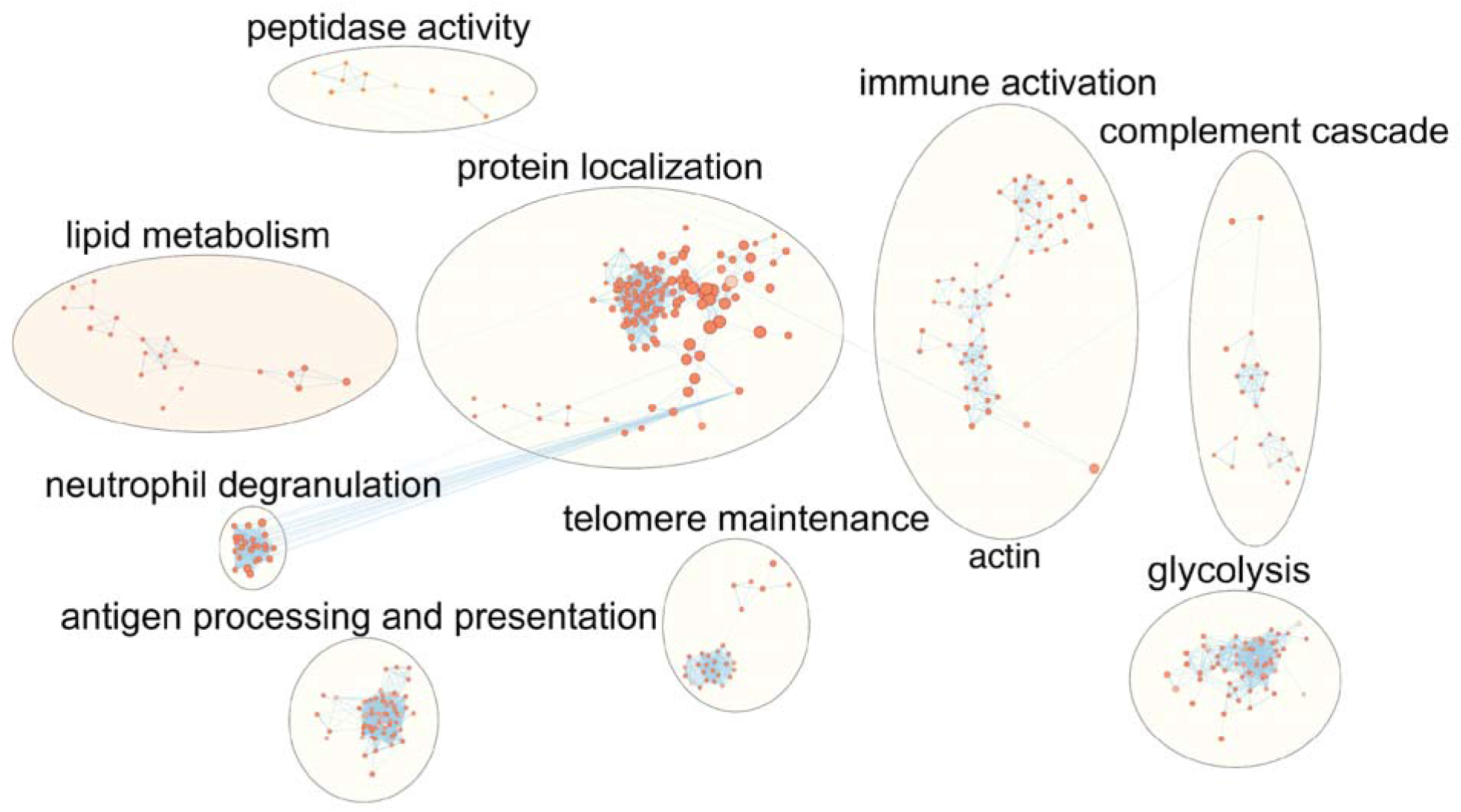
Proteomic analysis identifies altered lipid metabolism pathways in the PD appendix. Pathway enrichment analysis of proteomic changes in the PD appendix relative to controls (n=3 PD, 3 controls). Pathway analysis of quantitative proteomic data was performed using g:Profiler. Nodes are pathways altered in the PD appendix that were clustered into functionally similar networks by EnrichmentMap (nodes are *q*<0.05 pathways, hypergeometric test). Node size represents the number of genes in the pathway gene set, and edges connect pathways with similar gene sets (0.7 similarity cutoff). The lipid metabolism pathway network is highlighted in peach.

### Microbiome-driven bile acid changes in the appendix of PD patients and of the PD mouse model

In the appendix of PD patients and the mouse model of synucleinopathy, we observed changes affecting lipid metabolism and changes in microbiota that generate hydrophobic secondary bile acids (*Clostridium* cluster *XI*) and in those that are bile acid-resistant (*Burkholderiales*) (Wallner et al., 2019). Consequently, we examined whether there was a disturbance in bile acid levels in the appendix of PD patients. We quantified 15 bile acids in appendix tissue from PD patients and controls (n=15 and 12, respectively), examining primary and secondary bile acids (**Figure 4A**; Supplementary File 8). There was an 18.7-fold increase in LCA and a 5.6-fold increase in DCA in the appendix of PD patients relative to controls (*p*<0.05, robust linear regression adjusted for age, sex, and postmortem interval; **Figure 4**; Supplementary File 8). Similarly, group-level analysis examining bile acids with their taurine and glycine conjugates also showed a significant increase in LCA and DCA groups in the PD appendix (7.3-fold and 8.2-fold increase, respectively; *p*<0.05, robust linear regression; **Figure 4**). We did not observe any changes in primary bile acids or total bile acid levels in the PD appendix. Thus, the PD appendix exhibits elevated levels of hydrophobic, toxic bile acids that are produced by the microbiome.

**Figure 4:**
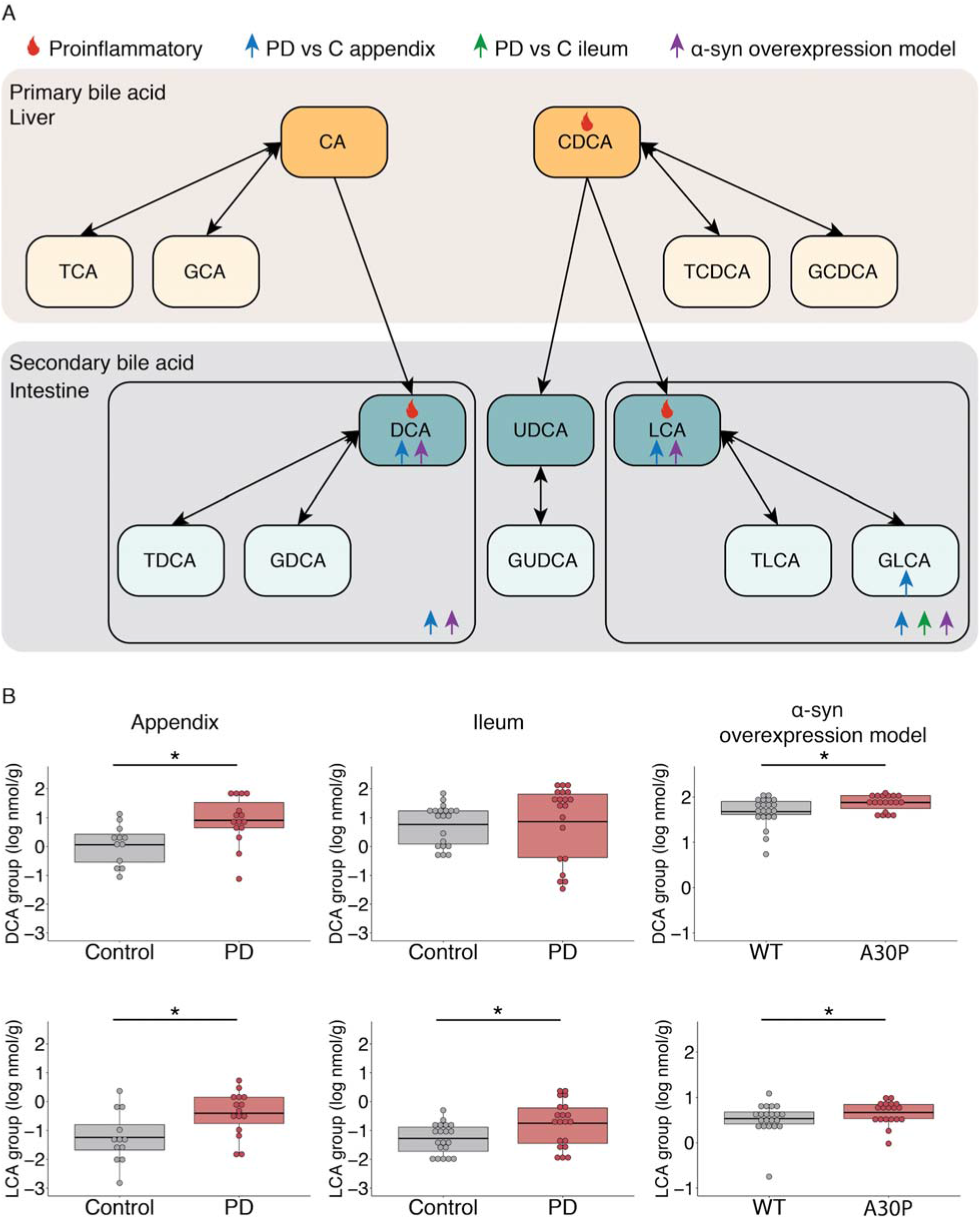
Increase in microbiome-derived secondary bile acids in the appendix of PD patients and synucleinopathy mouse model. Bile acid analysis was performed by liquid chromatography–mass spectrometry in the PD and control appendix (n=15 PD, 12 controls) and ileum (n=20 PD, 20 controls), as well as in the cecal patch of A30P α-syn overexpressing mice exposed to DSS or water). Bile acid changes were determined by robust linear regression, which for the human appendix and ileum data controlled for age, sex, and postmortem interval, and for the mouse cecal patch data controlled for sex and DSS exposure. (**A**) Illustration of the bile acid changes identified in this study and the bile acid pathway. Primary bile acids are generated in the liver and secondary bile acids are produced by the microbiome in the intestine. In the secondary bile acid section of the image, boxes highlight the DCA and LCA groups (DCA, LCA and their respective conjugates). Bile acids increased in the PD appendix or PD ileum, relative to controls, are marked by a blue and green arrow, respectively. Bile acids increased in the A30P α-syn mice, relative to wild-type mice, are marked by a purple arrow. The flame symbol denotes highly hydrophobic bile acids that have proinflammatory effects when elevated. (**B**) Secondary bile acid changes in the PD appendix, PD ileum, and in the synucleinopathy mouse model. The boxplot center line represents the mean, the lower and upper limits are the first and third quartiles (25^th^ and 75^th^ percentiles), and the whiskers are 1.5× the interquartile range. \**p*<0.05, robust linear regression

We next examined bile acid changes in the cecal patch of the mouse model of synucleinopathy and gut inflammation. As observed in the PD appendix, the cecal patch of mice that overexpress human A30P α-syn had an increase in LCA and DCA (1.3-fold and 1.4-fold increase, respectively; *p*<0.05, robust linear regression; **Figure 4**; Supplementary File 8). Likewise, group-level analysis of bile acids with their conjugates found an increase in the LCA and DCA groups in the cecal patch of A30P α-syn mice (1.3-fold and 1.4-fold increase, respectively; *p*<0.05, robust linear regression; **Figure 4**). Primary bile acids shared between the mice and humans were not altered by synucleinopathy. Furthermore, prior exposure to DSS colitis did not induce any bile acid changes in the cecal patch (Supplementary File 8). Therefore, a model of synucleinopathy mimics the secondary bile acid changes observed in the PD appendix.

The distal ileum is where primary bile acids are largely reabsorbed for return to the enterohepatic circulation; those that escape are subsequently converted to secondary bile acids by the gut microbiota in the large bowel (Wahlström et al., 2016). We examined whether there were bile acid changes in ileum tissue of PD patients relative to controls (n=20 PD and 20 controls). In the PD ileum, primary bile acids were unchanged. However, there was a significant increase in the LCA group in the PD ileum (3.6-fold increase; *p*<0.05, robust linear regression, adjusting for age, sex, and postmortem interval; **Figure 4**; Supplementary File 8). This signifies a prevalent accumulation of the most toxic bile acid in the PD gut.

### Bile-associated transcript and proteomic changes in the PD gut

Since we found a dysregulation of microbiome-derived bile acids in PD, we investigated whether PD patients had a differential expression of genes important for bile acid biosynthesis, signaling, and transport. In ileum tissue of PD patients and controls, we examined transcript levels of nuclear receptors that regulate bile acid and cholesterol homeostasis (FXR and LXR, respectively), the bile acid-sensing receptor (TGR5), transporters for bile reabsorption (ASBT, OSTα/OSTβ, FABP6), and transporters for cholesterol reabsorption and efflux (NPC1L1 and ABCG5/ABCG8, respectively). We also examined liver tissue of PD patients and controls for transcript levels of these genes or their liver-specific functional equivalents (NTCP and FABP1 for bile acid transport), as well as the rate-limiting enzymes for bile acid biosynthesis in the liver (CYP7A1 and CYP27A1). We found that the PD ileum had significantly elevated levels of gene transcripts involved in cholesterol homeostasis and transport: *NR1H2, NR1H3*, *NPC1L1*, *ABCG5*, and *ABCG8* (*p*<0.05, one-way ANOVA test; **Figure 5A**; Supplementary File 9). In the liver, we did not observe changes in bile acid-related transcripts (**Figure 5A**; Supplementary File 9).

**Figure 5:**
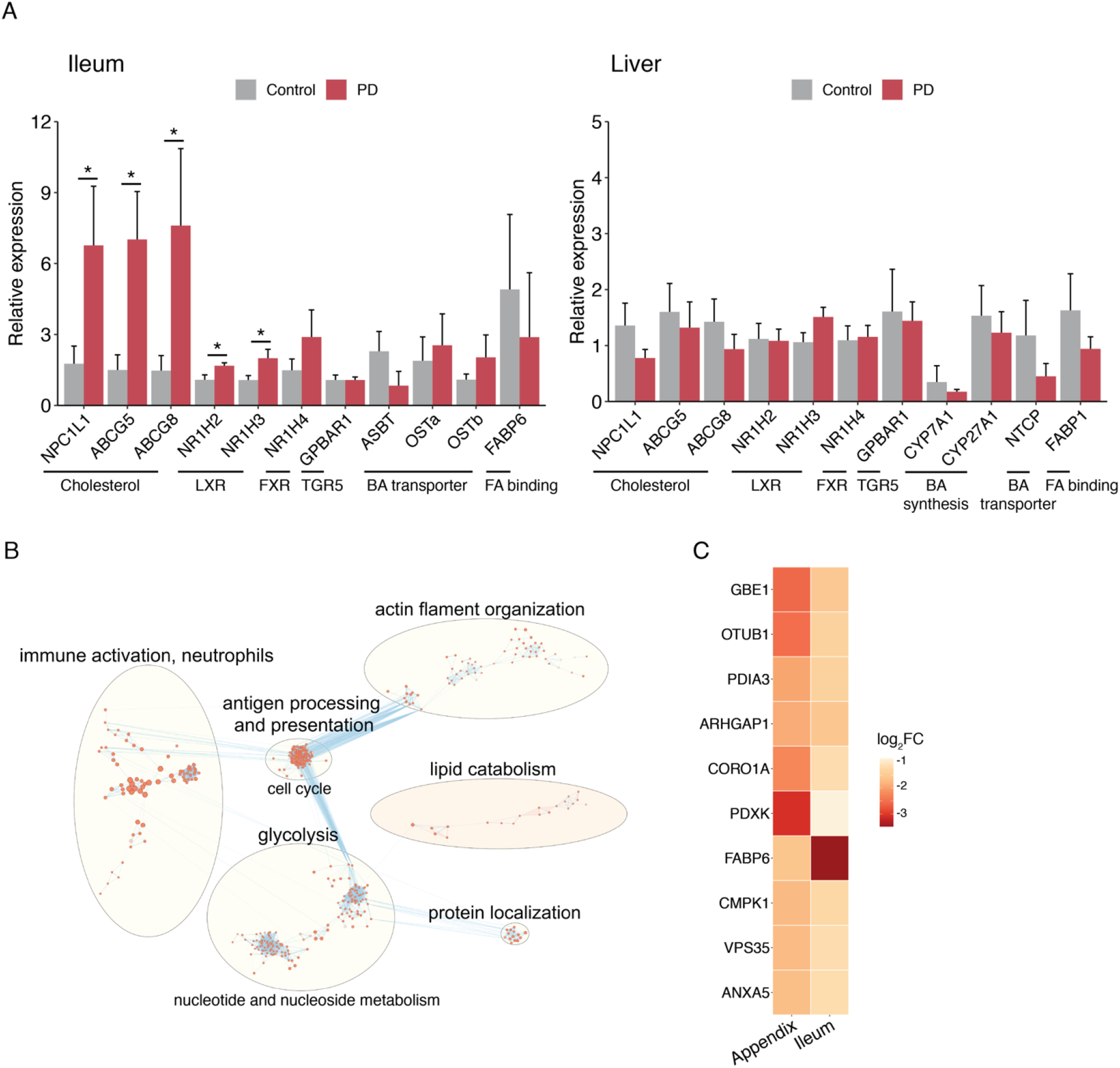
Dysfunctional cholesterol and lipid metabolism in the PD ileum. (**A**) Transcript levels of genes in the ileum (left panel) and liver (right panel) affecting the abundance of cholesterol and bile in the enterohepatic circulation. Transcript levels of genes affecting cholesterol and bile acid homeostasis (*NR1H2*, *NR1H3*, *NR1H4*, *GPBAR1*) and their transport and reabsorption into the enterohepatic circulation (*NPC1L1*, *ABCG5*, *ABCG8*, *ASBT, OST*α*, OST*β*, FABP6*) were examined in the ileum. In the liver, transcript levels of these genes or the equivalent bile acid transporters (*NTCP*, *FABP1*) were examined, as well as the rate-limiting enzymes for bile acid production (*CYP7A1*, *CYP27A1*). Transcript levels were analyzed by qPCR and normalized to housekeeping genes (villin1, β-actin). Relative expression ± s.e.m in the ileum (n=8 PD, 6 controls) and liver (n=6 PD and 6 controls). \**p*<0.05, one-way ANOVA. (**B**) Pathway analysis of proteomic changes in the PD ileum. Quantitative proteomic analysis in PD and control ileum was performed by mass spectrometry and changes in proteomic pathways were determined using g:Profiler (n=4 PD, 4 controls). Nodes are pathways altered in the PD ileum that were clustered into functionally similar networks by EnrichmentMap (nodes are *q*<0.05 pathways, hypergeometric test). Node size represents the number of genes in the pathway gene set, and edges connect pathways with similar gene sets (0.7 similarity cutoff). The lipid catabolism pathway network is highlighted in peach. (**C**) Top 10 proteins that were most consistently altered in the PD appendix and ileum. Heatmap showing the proteins ranked as the most consistently disrupted in the PD appendix and ileum, as determined by a robust ranking algorithm. Log fold change is shown, and red signifies greater disruption in PD.

We performed a human proteomic analysis to identify biological pathways altered in the ileum of PD patients relative to controls. The PD ileum had a decrease in lipid metabolism, as was observed in the PD appendix (*q*<0.05 pathways, hypergeometric test; **Figure 5B**). In the PD ileum, there was also a dysregulation of antigen processing and presentation, immune activation, glycolysis, and actin filament organization (*q*<0.05 pathways, hypergeometric test; **Figure 5B**). We next determined the proteins most consistently altered in the PD ileum and appendix using a robust ranking algorithm (Kolde et al., 2012). We found that both the PD ileum and appendix had a strong decrease of fatty acid binding protein 6 (FABP6), the intracellular bile acid transporter involved in returning bile acids to the enterohepatic circulation (**Figure 5C**). The PD ileum and appendix also had a loss of pyridoxal kinase (PDXK), which is essential for vitamin B_6_ bioactivity (Parra et al., 2018), and of vacuolar protein sorting 35 ortholog (VPS35), which is a PD risk factor that affects neurodegeneration (Chen et al., 2019b). Thus, the human transcript and proteomic changes in the PD gut are consistent with a disruption in bile acid control, including alterations in mediators of cholesterol homeostasis and lipid metabolism.

### Common blood markers are associated with PD in PPMI data

Given the increase in toxic secondary bile acids found in the PD gut, we investigated whether we could detect changes in markers of liver and gallbladder function in PD. We analyzed blood test results from the highly clinically characterized individuals in the Parkinson’s Progression Markers Initiative (PPMI). This initiative includes a *de novo* PD cohort who, at enrollment, had been diagnosed with PD for less than two years and were not taking PD medications. In this cohort, we examined five commonly used markers of liver abnormalities: alanine transaminase (ALT), aspartate aminotransferase (AST), alkaline phosphatase (ALP), albumin, and bilirubin. Though the mean values for each test were within the normal range, we found that *de novo* PD patients had a significant increase in ALP, bilirubin, and albumin, as well as a significant decrease in AST and ALT, relative to controls (*p*<0.05, robust linear regression adjusting for age, sex, and anemia measures; n=475 PD, 231 controls; **Figure 6A**; Supplementary File 10). *De novo* PD patients also had a higher prevalence of individuals with AST/ALT ratios above 1 (De Ritis ratio), a sign of liver injury (*p*<0.05, chi-squared test). Similarly, a cohort of PD patients with a genetic risk for the disease, carriers of a *LRRK2*, *GBA*, or *SNCA* mutation, had an increase in ALP and a decrease in AST and ALT (*p*<0.001, robust linear regression; n=396 PD, 619 controls; **Figure 6B**). Thus, newly diagnosed PD patients show notable changes in markers of biliary tract dysfunction.

**Figure 6:**
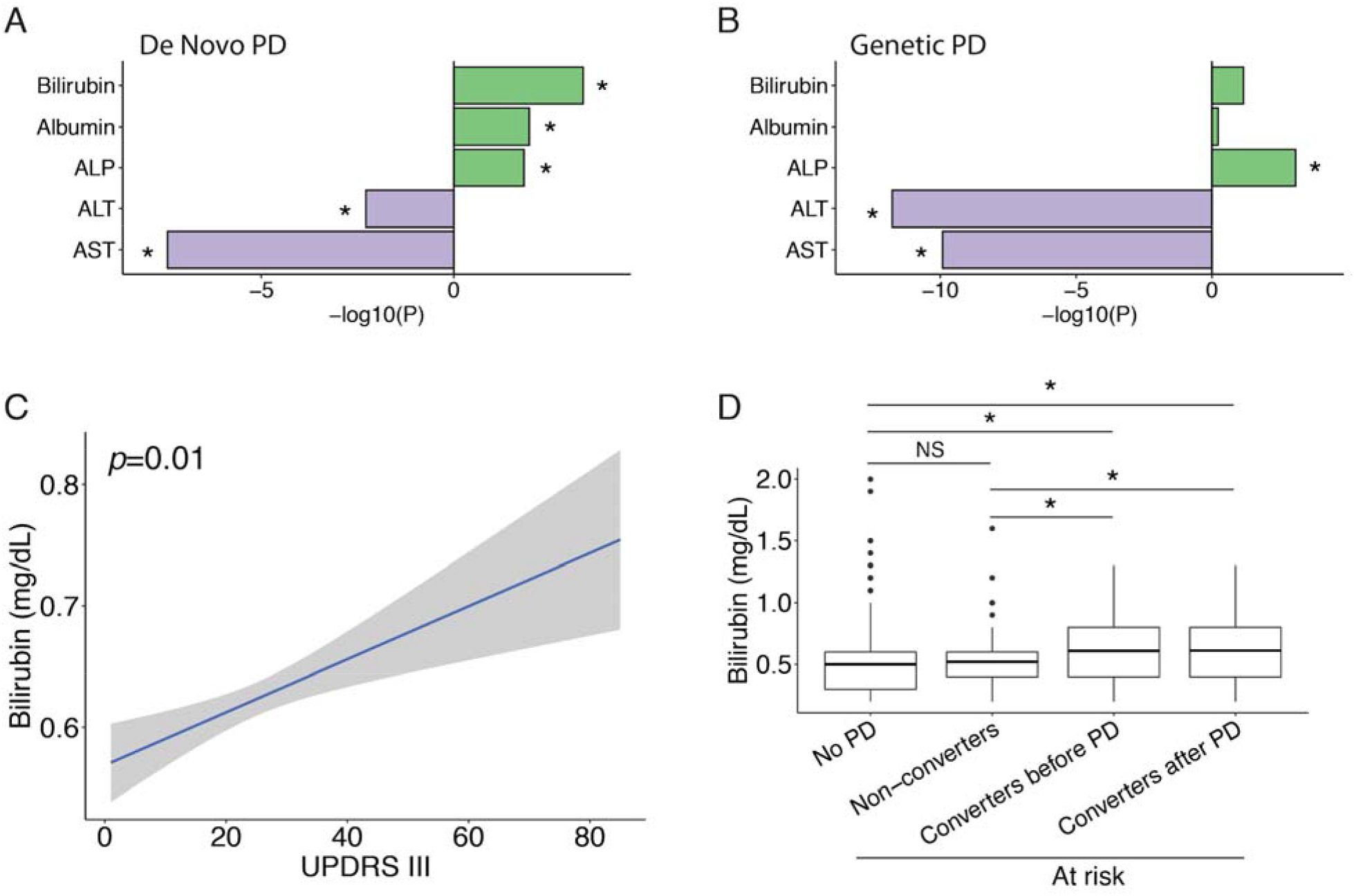
PD patients have biliary abnormalities that correspond to motor symptom severity and occur prior to PD diagnosis. Blood markers of liver and gallbladder function assessed in PD patients were bilirubin, albumin, ALP, ALT, and AST. Changes in blood markers were examined in newly diagnosed PD patients (*de novo* PD cohort, n=475 PD, 231 controls) and PD patients with a genetic risk factor (carriers of a mutation in *SNCA*, *LRRK2*, or *GBA*, n=396 PD, 619 controls) from the PPMI study. (**A, B**) Changes in common blood markers of liver and gallbladder disease in newly diagnosed PD patients (*de novo* PD, **A**) and with genetic risk for PD (genetic PD, **B**), compared with controls. Signed log p refers to the significance of blood marker change, with sign corresponding to the direction of the change in PD; green is increased, and purple is decreased in PD. \**p*<0.05, by robust linear regression, adjusting for age at the time of blood test, sex, and anemia measures (red blood cells, hemoglobin and hematocrit). (**C**) Bilirubin changes correlated to severity of PD motor symptoms. PD motor symptoms in *de novo* PD patients were determined by UPDRS score III. *p*=0.01 signifies that bilirubin levels increase with the severity of PD motor symptoms, by linear mixed-effects model, adjusting for disease duration, age of PD onset, sex, anemia measures, as well as UPDRS III score and disease duration interaction. **(D)** Bilirubin levels prior to PD diagnosis in individuals at risk of developing PD. Changes in blood markers were examined in the prodromal PD cohort, consisting of individuals with RBD or hyposmia (prodromal features of PD). Comparison of bilirubin levels in prodromal PD patients that will later be diagnosed with PD (n=36 converters) and those that will not (n=102 non-converters), relative to controls (n=561). Blood markers were assessed at the date of enrollment into the PPMI, prior to any PD diagnosis; and for converters we also examined bilirubin levels after PD diagnosis. \**p*<0.05 and NS=not significant, by robust linear regression, adjusted for age at the time of blood draw, sex, and anemia measures. The boxplot center line represents the mean, the lower and upper limits are the first and third quartiles (25^th^ and 75^th^ percentiles), and the whiskers are 1.5× the interquartile range.

We next examined whether changes in these blood markers were associated with the severity of PD motor symptoms, and whether these changes were predictive of PD conversion risk. We found that bilirubin levels positively correlated with PD motor symptom severity (n=472 *de novo* PD patients, UPDRS part III; *p*<0.05, linear mixed-effects model; **Figure 6C**; Supplementary File 10). No motor symptom associations were observed for the other blood markers. To further investigate these changes, we analyzed a PPMI cohort of individuals who have prodromal signs of PD (hyposmia or REM sleep behavior disorder) and are consequently at risk of developing PD. We analyzed bilirubin levels in subjects before PD diagnosis. Prior to PD diagnosis, individuals that would later be diagnosed with PD had elevated bilirubin levels compared to individuals who did not convert to PD and to controls (*p*<0.01, robust linear regression adjusted for age, sex, and anemia measures; n=36 PD converters, 102 PD non-converters, 561 controls; **Figure 6D**; Supplementary File 10). In addition, bilirubin levels remained significantly elevated after PD diagnosis relative to non-converters and to controls (*p*<0.05, robust linear regression; **Figure 6D**). Therefore, PD patients have biliary abnormalities that occur prior to PD diagnosis.

## Discussion

Our metatranscriptomic analysis revealed significant differences in the appendix microbiome of PD patients, which are closely related to bile acid dysregulation in the gut. First, in the PD appendix we found an increase in microbiota that generate hydrophobic secondary bile acids (*Clostridium* cluster *XI*) and that are bile acid-resistant (*Burkholderiales*), and several of these changes were recapitulated in a mouse model of synucleinopathy. Next, we analyzed bile acids in human appendix and ileum, as well as in the cecal patch of the synucleinopathy model and found a prominent increase in the hydrophobic bile acids LCA and DCA. Proteomic and transcript analysis supported a dysregulation of lipid metabolism and cholesterol homeostasis in the PD gut. Finally, an analysis of PD blood showed that markers of biliary tract dysfunction were present in both *de novo* and genetic forms of PD, and that bilirubin was elevated in at-risk individuals before a diagnosis of PD. In sum, our findings provide a novel look into the appendix microbiome in PD and demonstrate microbially-mediated bile acid disturbances in PD (Figure 7 and S6). Our finding of biliary alterations in the early stage of PD draws attention to the enterohepatic system, which has previously been overlooked in PD, but could hold therapeutic potential.

**Figure 7:**
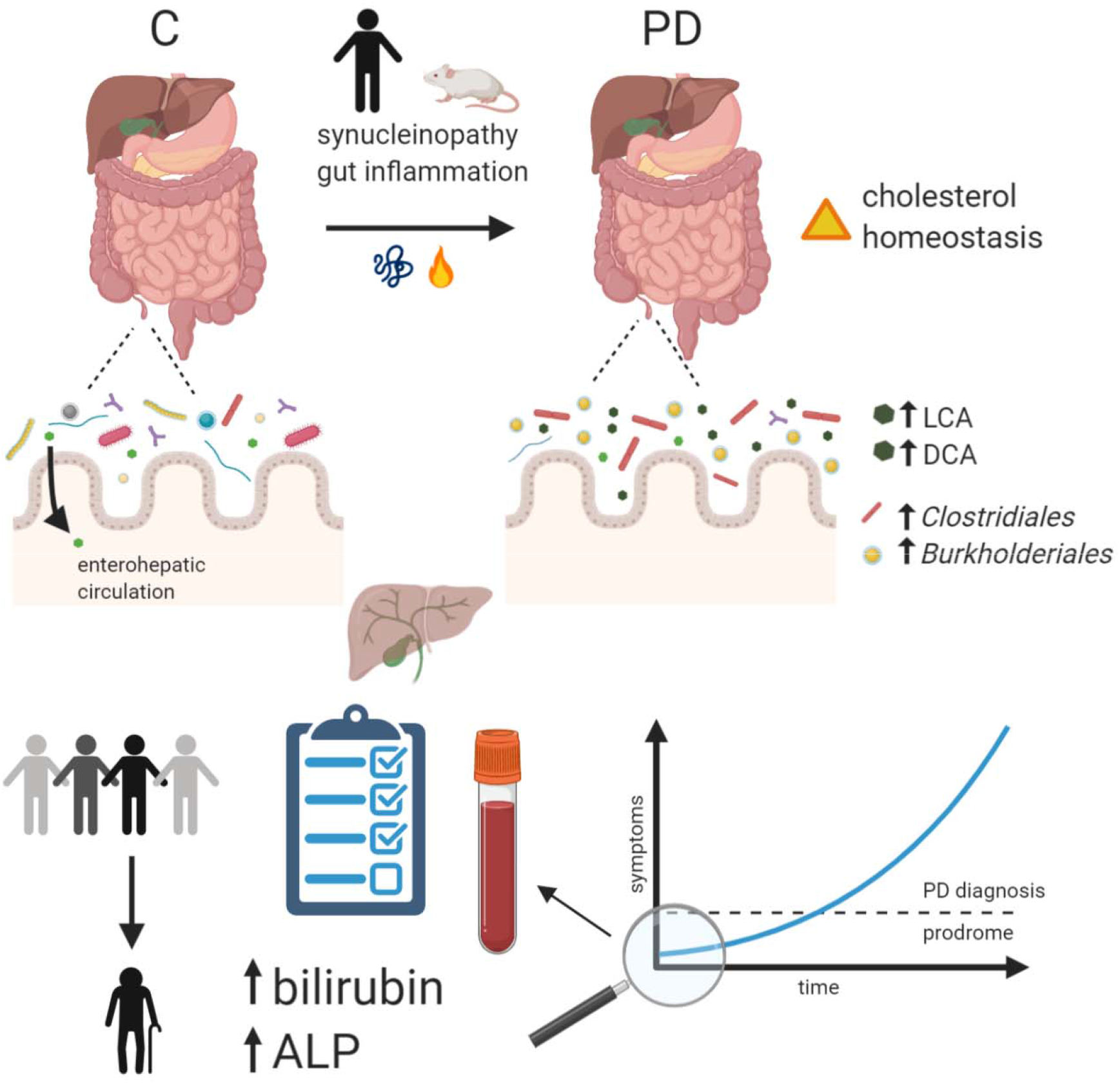
Graphical summary of microbiome and biliary changes in PD. Microbial dysbiosis and an accompanying increase in microbiome-derived toxic secondary bile acids are prevalent in the PD appendix. Synucleinopathy in mice recapitulates microbiome changes relevant to PD, some of which are amplified by gut inflammation. Microbiome changes in PD included significant increases in *Clostridiales* and *Burkholderiales,* which are involved in the conversion of primary bile acids to secondary bile acids. Accordingly, elevated secondary bile acids, LCA and DCA, were seen in PD and in the synucleinopathy model. Transcript and proteomic analyses of the PD gut showed further disruptions in cholesterol metabolism and transport. Bilirubin and ALP were elevated in newly diagnosed PD patients relative to controls. Bilirubin increases in PD correlated to the severity of motor symptoms. Moreover, in a prodromal cohort bilirubin was increased in individuals who would later convert to PD relative to controls and to individuals who did not convert to PD.

We found significant microbiome differences in the PD appendix, specifically, an increase in *Peptostreptococcaceae* (*Clostridium* cluster *XI*)*, Burkholderiales*, and *Lachnospiraceae* (which includes several genera in *Clostridium cluster XIVa*), as well as a decrease in *Methanobacteriales*. Several studies of the fecal microbiome have found dysbiosis in PD (Bedarf et al., 2017; Heinzel et al., 2020; Scheperjans et al., 2015; Weis et al., 2019); however, this study is the first to investigate the appendix microbiome in PD. Importantly, microbiome profiling was performed using a metatranscriptomic approach, which, unlike the prior 16S rRNA and metagenomic studies, has the advantage of capturing only living, active species and gives insight into functional mechanisms occurring during dysbiosis. Microbiome shifts in the appendix are particularly relevant given the role of the appendix in immunosurveillance and in affecting the microbiome in other intestinal regions (Killinger and Labrie, 2019). In addition, the microbial community we identified in our study of postmortem tissues was consistent with that of histologically normal, surgically-isolated human appendixes (Guinane et al., 2013; Jackson et al., 2014; Peeters et al., 2019). Differences between the appendix and fecal microbiomes limit comparison with previous studies of the PD microbiome; however, an increase in *Clostridiales cluster XI* in PD stool was observed by 16S rRNA analysis (Weis et al., 2019). Further, our findings are strengthened by the recapitulation of these changes in an α-syn overexpressing mouse model of PD, and thus it is unlikely that the microbiome changes were merely due to confounding environmental factors in the human cases. Our microbiome analysis does not distinguish between microbiome changes causing PD from those that are consequences of the disease, but our analysis of the synucleinopathy mouse model supports the relevance of the microbial changes to PD pathobiology.

Though it is established that the microbiome plays a role in human health and disease, the mechanisms by which microbiome changes could contribute to PD are not well understood. Some have proposed that the production of bacterial metabolites modulates inflammation and immune response in the brain (Erny et al., 2015; Fung et al., 2017; Sampson et al., 2016). Microbiome-produced short chain fatty acids were shown to enhance neuroinflammation in the brain and behavioral motor deficits in a mouse model of synucleinopathy (Sampson et al., 2016). In addition, microbiome alterations are associated with ulcerative colitis and Crohn’s disease (inflammatory bowel diseases, IBD), which have an increased co-occurrence with PD (Hui et al., 2018; Villumsen et al., 2019). The microbiome can also affect autophagy, another cellular process implicated in PD (Donohoe et al., 2011; Lin et al., 2014; Xilouri et al., 2016). Our results implicate another pathway: microbial bile acid metabolism in PD.

Bacterial species belonging to *Clostridium cluster XI and Cluster XIVa,* which include species in the families *Peptostrococcaceae and Lachnospiraceae,* are responsible for the conversion of primary bile acids to the more hydrophobic secondary bile acids in the large intestine; these were elevated in the PD appendix and in the cecal patch of our experimental mouse model. In both the PD appendix and α-syn overexpressing mouse model, there was also an increase in *Burkholderiales*. *Burkholderia* is a bacterial genus with a broad environmental distribution that is recognized to be an opportunistic, antibiotic-resistant pathogen. It can cause severe inflammation in immunocompromised individuals (Chiarini et al., 2006) and produce kynurenine and quinolinate (Kaur et al., 2019), which are proinflammatory metabolites associated with symptom severity in PD (Heilman et al., 2020). *Burkholderia* can also infect the brain via propagation in nerves (St John et al., 2016) or via peripheral immune cell entry across the blood-brain barrier (Hsueh et al., 2018). Of particular interest, *Burkholderia* species encode the rate-limiting enzyme for secondary bile acid synthesis (bile-acid dehydratase, baiE) (Wallner et al., 2019). Thus, the PD appendix microbiome has an enrichment of microbiota that metabolize bile acids.

In line with the microbiota shifts seen in the PD appendix, we found an increase in the secondary bile acids LCA and DCA in both the appendix and ileum of PD patients. LCA and DCA are highly hydrophobic and their increase has been long recognized to have direct cytotoxic effects (Chen et al., 2019a; Pavlidis et al., 2015). An accumulation of secondary bile acids can disturb cell membranes and epithelial barrier integrity leading to the generation of reactive oxidative species and induction of apoptosis (Hegyi et al., 2018). Furthermore, LCA and DCA alter host inflammatory responses and may contribute to PD pathogenesis through such a mechanism. DCA activates NF-κB, a transcription factor that is a key regulator of proinflammatory responses (Huo et al., 2011; Liu et al., 2017). Secondary bile acids, particularly LCA, control the differentiation and activation of regulatory T (T_reg_), T helper, and natural killer T (NKT) cells (Hang et al., 2019; Ma et al., 2018; Song et al., 2020), and there is evidence for T cell involvement in early PD (Lindestam Arlehamn et al., 2020). Gut inflammation has been shown to trigger pathogenic α-syn accumulation (Kishimoto et al., 2019; Stolzenberg et al., 2017). Since bile acids can act as signaling molecules that modulate immune response (Duboc et al., 2013; Guo et al., 2016; Hang et al., 2019; Song et al., 2020), the microbial-driven bile acid changes may be a response to α-syn aggregation in the PD gut. In addition, the membrane-damaging and inflammatory effects of raised hydrophobic bile acids in PD could propel the accumulation of pathological α-syn aggregates, which could potentially propagate from gut to brain. Further work in *in vivo* models will be necessary to elucidate the effects of secondary bile acids on PD pathology.

Our study gives several insights into the potential causes behind the changes in the microbiome and bile acid composition in PD. Though we cannot exclude constipation as a contributing factor, it is unlikely, because LCA and DCA increase colonic peristalsis and accelerate fecal transit time (Alemi et al., 2013; Misawa et al., 2020). Also, these secondary bile acids are reduced in mice given fecal microbiome transplants from constipated donors (Ge et al., 2017). Rather, our mouse experiments point to synucleinopathy as a contributor to the observed changes in secondary bile acids in PD. As in PD, A30P α-syn overexpressing mice had a significant increase in both LCA and DCA. In PD patients, there was no evidence of an overproduction of primary bile acids because there was no significant increase in primary bile acids or total bile acid pool size in the PD ileum, and there was no transcript abnormalities in enzymes responsible for primary bile acid synthesis in the PD liver. However, PD patients may have impaired bile acid reuptake in the ileum, as indicated by the prominent decrease in FABP6, a protein responsible for efficient transport of primary bile acids through enterocytes for recirculation (Praslickova et al., 2012). Thus, disrupted bile acid transport in the ileum and increased bile-metabolizing bacteria in the large bowel could together contribute to the raised levels of secondary bile acids in PD. This may also contribute to the observed abnormalities in cholesterol homeostasis in the PD gut, as demonstrated by the transcriptional increase in cholesterol transporters and proteomic disruption of lipid metabolism pathways. It is also worth noting that changes in bile acids can in turn modulate the composition of the microbiome; bile acids exert strong selective pressure on the gut microbiota. Though this study does not delineate which changes occur first in PD, the bi-directional relationship between the microbiome and bile acid composition creates a system which, once triggered, may lead to a self-reinforced condition of dysbiosis, peripheral inflammation, and α-syn aggregation.

Disruption of bile acid homeostasis has been linked to liver and gallbladder disease. In our analysis of blood markers, we found newly diagnosed, idiopathic PD patients had indicators of liver and gallbladder dysfunction, with an increase in bilirubin, ALP and in the ALT:AST ratio. Likewise, ALP levels were increased in PD patients with genetic risk for the disease (carriers of a *SNCA*, *LRRK2*, or *GBA* risk variant). Notably, the mean values of these markers in PD patients still fell within normal ranges, signifying that PD patients overall are not suffering from clinical liver or gallbladder disease at the time of testing. However, the significant differences in PD patients could indicate sub-clinical abnormalities that nevertheless provide meaningful clues about PD pathobiology. Indeed, liver injury may have a role in PD. Numerous epidemiological studies have shown that chronic liver diseases are associated with a greater risk of developing PD (Burkhard et al., 2003; Lin et al., 2019; Pakpoor et al., 2017). Elevated secondary bile acids in the enterohepatic system can induce liver injury; both DCA and LCA are hepatotoxic, and LCA administration in animals is a common model of intrahepatic cholestasis (blocked bile ducts) (Song et al., 2011). Secondary bile acids also contribute to gallstone disease (Berr et al., 1996; Di Donato et al., 1986; Marcus and Heaton, 1988). Both liver dysfunction and gallstone disease can lead to elevated blood levels of bilirubin, and, interestingly, we found that bilirubin levels were increased in PD and associated with the severity of motor dysfunction in patients (UPDRS part III score). Bilirubin changes were also found to precede PD diagnosis, pointing to biliary abnormalities early in PD. Thus, the increase in secondary bile acids observed in PD may cause hepatobiliary abnormalities that are detectable before PD diagnosis.

Targeting the appendix microbiome and bile acids may be an innovative approach for future therapeutics. Microbiome transplantation effectively restores the commensal microbiome in intestinal infections (Mullish et al., 2019) and has been proposed for the treatment of PD. Since the appendix contains a thick, shielded biofilm matrix in which bacteria reside (Killinger and Labrie, 2019; Palestrant et al., 2004), this could be a strategic location to administer a long-lasting transplant. Experimental models indicate that the appendix can influence the microbiome in the large intestine, and routine shedding of biofilm fragments from the appendix may repopulate the intestine with the healthy, transplanted microbiome (Donaldson et al., 2016; Masahata et al., 2014). Our results suggest that bacteria that withstand bile acids and promote DCA and LCA conjugation to its less toxic derivatives may be most therapeutic. In addition, preventing the damaging effects of LCA and DCA with the hydrophilic, anti-inflammatory bile acid UDCA could benefit PD patients. UDCA has been shown to counteract the effects of the hydrophobic secondary bile acids in the liver and gallbladder (Rodrigues et al., 1998; Salen et al., 1980). UDCA also has neuroprotective effects in PD models (Mortiboys et al., 2015) and is currently being tested in clinical trials for PD (Sathe et al., 2020). In addition to therapeutic applications, measures of blood markers of liver abnormalities may be useful for PD prognosis in at-risk individuals. In particular, a longitudinal shift to increased bilirubin may signal a risk of phenoconversion to PD.

In sum, our results point to specific changes in the microbiome of PD patients that suggest biliary dysfunction as a previously unexplored mechanism involved in PD. Though further investigation is needed, bile acids could play a key role at the intersection of microbiome dysbiosis, inflammation, and α-syn misfolding. Considering the relative accessibility of the GI tract and existing therapies for bile acid-related disorders, targeting microbial-derived secondary bile acids may be a new avenue for earlier diagnosis and alleviation of PD symptoms.

## Acknowledgements

We thank the Van Andel Institute Genomics and Bioinformatics and Biostatistics Cores and Michigan State University Genomics Core. We thank the Oregon Brain Bank, Parkinson’s UK Brain Bank, and the NIH NeuroBioBank for the tissue provided. This work was supported by a Farmer Family Foundation grant award to PB, with LB, AP, and VL as co-investigators. VL is supported by grants from the Department of Defense (W81XWH1810512), the National Institute of Neurological Disorders and Stroke (1R21NS112614-01, 1R01NS113894-01A1, 1R01NS114409-01A1), and a Gibby & Friends vs. Parky Award.

Data used in the preparation of this article were obtained from the Parkinson’s Progression Markers Initiative (PPMI) database (www.ppmi-info.org/data). For up-to-date information on the study, visit www.ppmi-info.org. PPMI – a public-private partnership – is funded by the Michael J. Fox Foundation for Parkinson’s Research and funding partners, including Abbvie, Allergan, Amathus Therapeutics, Avid Radiopharmaceuticals, Biogen, BioLegend, Bristol-Myers Squibb, Celgene, Denali, GE Healthcare, Genentech, GlaxoSmithKline, Golub Capital, Handl Therapeutics, Insitro, Janssen Neuroscience, Lilly, Lundbeck, Merck, Meso Scale Discovery, Pfizer, Piramal, Prevail Therapeutics, Roche, Sanofi Genzyme, Servier, Takeda, Teva, UCB, Verily, and Voyager Therapeutics.

## Author contributions

PL contributed to the experimental design and the analysis of the microbiome, bile acids, proteomic and PPMI data. BK generated the microbiome libraries in human and mouse. IB analyzed the metatranscriptomic data. EE and NL contributed to the qPCR analysis. AY and SG generated the bile acid data. JL, MS, and IV were involved in the proteomic sample preparations and mass spectrometry. MB provided the tissue samples from the synucleinopathy and DSS model. AP, PB, and LB were involved in experimental design. VL was involved with study design and overseeing the experiments. The manuscript was written by VL and EE and commented on by all authors.

## Data Availability

All sequencing data used in this study are available from the NCBI Gene Expression Omnibus (GEO) database under the accession number GSE135743 and GSE156647. The mass spectrometry proteomics data of human ileum has been deposited to the ProteomeXchange Consortium via the PRIDE (Perez-Riverol et al., 2019) partner repository with the dataset identifier PXD020988. The human appendix proteomic data was obtained from PXD015079. Custom code for metagenomic and 16S rRNA sequencing analysis is available at https://github.com/lipeipei0611/bile_acid/.

## Competing interests

MB is a full-time employee at Roche and may additionally hold Roche stock/stock options. P.B. has received commercial support as a consultant from Axial Biotherapeutics, Calico, CuraSen, Fujifilm-Cellular Dynamics International, IOS Press Partners, LifeSci Capital LLC, Lundbeck A/S, Idorsia and Living Cell Technologies LTD. He has received commercial support for grants/research from Lundbeck A/S and Roche. He has ownership interests in Acousort AB and Axial Biotherapeutics and is on the steering committee of the NILO-PD trial. No other authors have conflicts of interest.

## Materials and Methods

### Human Tissue Samples

Human appendix, ileum, and liver tissue from PD patients and controls was obtained from the Oregon Brain Bank. For each individual, we had information on demographics (age, sex), tissue quality (postmortem interval), and pathological staging (Supplementary File 1). PD cases were pathologically confirmed to have brain Lewy body pathology, and control individuals had no brain Lewy body pathology. All human postmortem tissue work had the approval from the ethics committee of the Van Andel Institute (IRB #15025).

### Metatranscriptomic analysis of PD appendix microbiome

To profile functional microbiome changes in the PD appendix, we performed a metatranscriptomic analysis of the appendix of 12 PD and 16 controls. Frozen appendix tissue (∼20 mg) was homogenized using a Covaris cryoPREP pulverizer and then in 1 mL of TRIzol (Life Technologies) with a ceramic bead-based homogenizer (Precellys, Bertin Instruments). Total RNA was isolated according to the TRIzol manufacturer’s instructions, treated with RNase-free DNase I (Qiagen) at room temperature for 30 min, followed by clean-up with the RNeasy Mini Kit (Qiagen). Total RNA yield and quality was determined using a NanoDrop ND-1000 (Thermo Fisher Scientific) and an Agilent Bioanalyzer 2100 system (Agilent Technologies). Libraries were prepared by the Van Andel Genomics Core from 300 ng of total RNA using the KAPA RNA HyperPrep Kit with RiboseErase (v1.16; Kapa Biosystems). RNA was sheared to 300-400 bp. Prior to PCR amplification, cDNA fragments were ligated to NEXTflex dual adapters (Bioo Scientific). The quality and quantity of the finished libraries were assessed using a combination of Agilent DNA High Sensitivity chip (Agilent Technologies, Inc.), QuantiFluor dsDNA System (Promega Corp.), and Kapa Illumina Library Quantification qPCR assays (Kapa Biosystems). Individually indexed libraries were pooled, and 100-bp, single-end sequencing was performed on an Illumina NovaSeq6000 sequencer using an S1 100 cycle kit (Illumina Inc.), with all libraries run on a single lane to return an average depth of 37 million reads per library. Base calling was done by Illumina RTA3, and output of NCS was demultiplexed and converted to FastQ format with Illumina Bcl2fastq v1.9.0.

Preprocessing of the metatranscriptomic data involved the removal of sequencing adapters and low-quality bases from sequencing reads using Trim Galore (v0.5.0). The transcriptomic data was aligned to the human genome (GRCh38/hg38) with the twopassMode basic algorithm in STAR (v2.5.2b) (Dobin et al., 2013). Reads that did not align to the human genome (STAR option outReadsUnmapped) were then input to MetaPhlAn2 (v2.7.7) (Segata et al., 2012), which gives kingdom to species-level resolution for bacteria, archaea, eukaryotes, and viruses within the database (db_v20). We then performed functional profiling of the microbial community for the same non-human reads using HUMAnN2 (v0.11.1) (Franzosa et al., 2018) with the UniRef90 database. Pathway abundance data was normalized to counts per million using inbuilt HUMAnN2 functionality. To test for differential taxa abundance between PD patients and control, proportional microbial compositional data from MetaPhlAn2 was imported into R (v3.6) and converted back to counts for all taxa-level ids (features). To perform statistical analysis, we used the cumulative sum scaling normalization and the zero-inflated gaussian mixture model from the Bioconductor package metagenomeSeq (v1.28.0) (Paulson et al., 2013). Feature and pathway abundance data were examined using the fitZig function to determine microorganisms and pathways related to PD, adjusting for age, sex, postmortem interval, and RIN. P-values were derived from the empirical Bayes moderated F-statistic and adjusted for multiple testing correction using the Benjamini-Hochberg method, with *q*<0.05 set as the threshold for statistical significance.

### Microbiome analysis in a mouse model of synucleinopathy and gut inflammation

We examined microbiome changes by 16S rRNA sequencing of the V3-V4 hypervariable region in a mouse model of synucleinopathy as well as in response to gut inflammation. Microbiome changes were profiled in wild-type mice and in the synucleinopathy model: hemizygous Tg(Thy1-SNCA*A30P)18Pjk mice (A30P α-syn), which overexpress human α-syn with the A30P mutation under the neuron-selective Thy1 promoter (Kahle et al., 2000). A30P α-syn mice have been maintained on a C57BL/6 background for over 10 generations. Gut inflammation was induced using dextran sodium sulfate (DSS), a widely used model of ulcerative colitis (Chassaing et al., 2014; Grathwohl et al., 2019). Adult wild-type and A30P α-syn mice (3-months of age) were exposed to a chronic DSS protocol that began at DSS concentration of 2.5% and increased to 4% over 4 cycles (+0.5% increment per cycle). One cycle consisted of 5 days of DSS, followed by 2 days of water (DSS: 160110, MP Biomedicals, LLC). The non-DSS colitis groups were administered normal drinking water. Mice were then given a 4-week long recovery period during which they received normal drinking water, followed by tissue harvest. Mice were anesthetized with pentobarbital and transcardially perfused with PBS before tissue collection. Cecal patch tissue was snap frozen and stored at –80°C until processing for DNA isolation. Approximately equal numbers of males and females were included in each genotype (wild-type, A30P α-syn) and treatment (water, DSS) group (Supplementary File 3). The animal experiments were endorsed by a Roche internal review board and approved by the local animal welfare authorities in Canton Basel-Stadt, Basel, Switzerland.

Frozen cecal patch samples were ground on liquid nitrogen into a fine powder. DNA was then isolated from the tissue powder using the Powersoil DNA isolation kit (Qiagen) according to manufacturer’s protocol and quality was determined by a NanoDrop 2000 spectrophotometer (Thermo Fisher Scientific). The 16S rRNA concentration in each sample was confirmed by qPCR using the forward primer: 5’-TCCTACGGGAGGCAGCAGT, and reverse primer: 5’-GGACTACCAGGGTATCTAATCCTGTT. Cecal patch samples were then processed for sequencing according to the Illumina’s 16S Metagenomic sequencing library protocol (doc. #15044223). Briefly, 16S rRNA was amplified from samples using forward primer: 5’-TCGTCGGCAGCGTCAGATGTGTATAAGAGACAGTCGTCGGCAGCGTCAGATGTGTATAAG AGACAGCCTACGGGNGGCWGCAG, and reverse primer: 5’-GTCTCGTGGGCTCGGAGATGTGTATAAGAGACAGGTCTCGTGGGCTCGGAGATGTGTATA AGAGACAGGACTACHVGGGTATCTAATCC. The 16S rRNA V3-V4 region amplicon (550 bp) was verified by gel electrophoresis. The amplicon was then purified using AMPureXP beads. The purified amplicon was indexed using Nextera XT index kit according to the manufacturer’s protocol. Sample libraries were purified with AMPure XP beads, verified with an Agilent 2100 Bioanalyzer, and quantified with a Qubit dsDNA HS kit on a Qubit 3.0 fluorometer (Thermo Fisher Scientific). Sample libraries were then pooled in equimolar amounts. The Michigan State University Genomics Core performed pooled library quality and quantification using a combination of the Qubit dsDNA HS kit, Advanced Analytical Fragment Analyzer High Sensitivity DNA NGS and Kapa Illumina Library Quantification qPCR assays. Sequencing was performed on an Illumina MiSeq v3 flow cell, loading a library concentration of 3 pM and a 30% spike-in of the Illumina PhiX control DNA library. Sequencing was performed in a 300 bp paired-end format on an Illumina MiSeq. Base calling was done by Illumina Real Time Analysis (RTA) v1.18.54, and the output of RTA was demultiplexed and converted to FastQ format with Illumina Bcl2fastq v2.19.1.

For microbiome data analysis, we first removed adapters and low-quality reads (Q<30) from the sequencing reads with Trim Galore (v0.4.4). QIIME2 was used to profile the microbial community (Bolyen et al., 2019). In QIIME2, reads were first denoised and dereplicated with the dada2 algorithm prior to being classified against the SILVA database (v132) with a clustering at 99% sequence identity criterion. For microbial pathway analysis, KEGG pathway abundances were estimated using the PICRUSt2 software (Langille et al., 2013). The taxa and KEGG abundance data were imported into R (v3.5.1). A zero-inflated gaussian mixture model from metagenomeSeq (v1.28.0) was used to determine microorganisms and pathways altered by synucleinopathy (A30P α-syn genotype), adjusting for sex and DSS exposure, and altered by DSS-mediated gut inflammation, adjusting for sex and genotype. To determine the differential responses of A30P α-syn mice to gut inflammation, we examined genotype and DSS exposure interactions, adjusting for sex, using a zero-inflated gaussian mixture model from metagenomeSeq. We then used the contrasts.fit function from limma to identify microbiota differences in genotypes in response to DSS exposure with the following contrasts matrix: (DSS - Water in A30P syn mice) - (DSS - Water in wild-type mice). P-values were adjusted for multiple testing correction using the Benjamini-Hochberg method, with *q*<0.05 set as the threshold for statistical significance.

### Microbiome comparison with histologically normal surgical appendix samples

To confirm that the postmortem appendix tissues used in our metatranscriptomic analysis had a similar microbiome composition to appendix samples from living individuals, we analyzed 16S rRNA sequencing data from three histologically normal, surgically-isolated appendix samples (accession ID: SRP035179, ages≥13) (Jackson et al., 2014). The 16S rRNA sequencing (Jackson et al., 2014). The 16S rRNA sequencing analysis for the surgically-isolated appendix samples was performed as described above. We then performed a Pearson’s correlation analysis to explore the similarity of microbiome composition in postmortem appendix tissue with normal surgically-isolated appendix tissues, examining microbial taxa at both the order and family level.

### Bile acid sample preparation and metabolite quantification

Liquid chromatography–mass spectrometry (LC-MS) was used to measure primary and secondary bile acids in the appendix, and ileum of PD patients and controls, and in the cecal patch of the mouse model of synucleinopathy and gut inflammation. Tissues (25 mg) were homogenized using a bead homogenizer at 5,500 rpm for 30 s in 300 µL of extraction solvent (85% ethanol and 15% phosphate-buffered saline solution). Samples were then sonicated at 4°C for 10 min. Proteins and other impurities were removed by centrifugation at 13000 × *g* for 30 min at 4°C. The supernatant was collected and 10 µL was loaded onto the Biocrates Bile Acid kit (Biocrates Life Sciences). Data were acquired using an Acquity I-class (Waters) coupled with a Xevo TQ-S mass spectrometer (Waters). All specimens were acquired in accordance with the protocol for the Biocrates Bile Acids kit. Bile acid concentrations (nmol per gram of tissue weight) were calculated utilizing the Biocrates MetIDQ software and TargetLynx (Waters). For group-level analyses, the concentration of the bile acid and its glycine and taurine conjugates were summed. For example, the LCA group is the sum of LCA, GLCA, and TLCA, and the DCA group is the sum of DCA, GDCA and TDCA. For all-primary, all-secondary, and total bile acid analyses, the respective bile acids and their conjugates were summed.

Bile acid data were normalized by log10 transformation, as previously described (Pan et al., 2017). Bile acid changes in the PD appendix and ileum and in the cecal patch of the mouse model of synucleinopathy and gut inflammation were determined by multivariate robust linear regression models with empirical Bayes from the limma (v3.30.13) statistical package (Ritchie et al., 2015). For the appendix and ileum, we determined bile acid changes in PD, adjusting for age, sex, and postmortem interval. For the mouse cecal patch, we determined bile acid changes induced by A30P α-syn overexpression, adjusting for sex and DSS exposure, and altered by DSS-mediated gut inflammation, adjusting for sex and genotype. To determine whether bile acids were differentially altered in A30P α-syn mice in response to gut inflammation, we examined genotype and DSS exposure interactions, adjusting for sex, in the limma robust linear regression model, followed by a contrasts.fit function with the following contrasts matrix: (DSS - Water in A30P syn mice) - (DSS - Water in wild-type mice). *P*-value <0.05 was the threshold for statistical significance.

### Mass spectrometry and proteomics analysis

Quantitative proteomic analysis was performed to determine host biological pathways altered in the PD appendix and ileum. For this analysis, we used existing proteomic data from the PD and control appendix (n=3 individuals/group; PXD015079) and generated new proteomic data for the PD and control ileum (n=4 individuals/group). Mass spectrometry for the appendix and ileum samples was performed using the same protocol by the Integrated Mass Spectrometry Unit at Michigan State University. The wet tissue weight of each sample (∼30 mg tissue/sample) was measured and 5-fold lysate buffer (20 mM Tris Base (pH 7.4), 150 mM NaCl, 1 mM EGTA, 1 mM EDTA, 5mM sodium pyrophosphate, 30 mM NaF, 1x Halt Protease Inhibitor Cocktail (Thermo Fisher Scientific)) was used to homogenize the tissue on ice with a tissue grinder (Tissue Master 125, Omni International). The homogenate was centrifuged at 18,407 × *g* for 10 min at 4°C and the supernatant was retained. Protein concentration in each sample was determined using a BCA assay (Pierce BCA Protein Assay, Thermo Fisher Scientific). Protein lysates (10 μg) were denatured using 25 mM ammonium bicarbonate/80% acetonitrile and incubated at 37°C for 3 h. The samples were dried and reconstituted in 50 μl of 25 mM ammonium bicarbonate/50% acetonitrile/trypsin/LysC solution (1:10 and 1:20 w/w trypsin:protein and LysC:protein respectively) and digested overnight at 37°C. The samples were dried and reconstituted in 50 μl of 25 mM ammonium bicarbonate/5% acetonitrile.

Samples were loaded onto an UltiMate 3000 UHPLC system with online desalting. Each sample (10 μl) was separated using a C18 EASY-Spray column (2 μm particles, 25 cm x 75 μm ID) and eluted using a 2 h acetonitrile gradient into a Q-Exactive HF-X mass spectrometer. Data dependent acquisition for Full MS was set using the following parameters: resolution 60,000 (at 200 m/z), AGC target 3e6, maximum IT 45 s, scan range 300 to 1500 m/z, dynamic exclusion 30 s. Fragment ion analysis was set with the following parameters: resolution 30,000 (at 200 m/z), AGC target 1e5, maximum IT 100 ms, TopN 20, isolation window 1.3 m/z, NCE at 28. Each sample was run in triplicate. The mass spectra from each technical replicate were searched against the Uniprot human database (filtered-proteome_3AUP000005650) using the LFQ method in Proteome Discoverer (v. 2.2.0.388, 2017) set as follows: at least 2 peptides (minimum length=6, minimum precursor mass=350 Da, maximum precursor mass 5000 Da), tolerance as set to 10 ppm for precursor ions and 0.02 Da for fragment ions (b and y ions only), dynamic modification was set for methionine oxidation (+15.995 Da) and N-terminus acetylation (+42.011 Da), target FDR (strict minimum value 0.01), Delta Cn minimum value 0.05). LFQ was calculated using the following parameters: ratio calculation set to pairwise ratio based, maximum allowed fold change 100, ANOVA (background based). The technical replicates from each biological sample were pooled to perform diagnosis comparisons, using a non-nested test. Proteins were quantified using the pairwise peptide ratio information from extracted peptide ion intensities. Only proteins with abundances recorded in at least 50% of samples were considered. Proteins with a log fold change between groups exceeding ± 0.2 were considered altered. Pathway analysis of proteins altered in the PD appendix and ileum was performed using g:Profiler (Raudvere et al., 2019), with networks determined by EnrichmentMap and clustered by AutoAnnotate in Cytoscape (v3.7.1) (Reimand et al., 2019). To identify the proteins that were most altered in both the PD appendix and ileum, we first determined the proteins that exhibited significant changes and had the same direction of change in both the PD appendix and ileum. These altered proteins were ranked by log fold change, ranking separately for appendix and ileum. We then determined the proteins most consistently altered in the PD appendix and ileum using the aggregateRanks function from the RobustRankAggreg package (v1.1) (Kolde et al., 2012).

### qPCR analysis of gene transcripts involved in bile acid and cholesterol homeostasis

We examined transcriptional changes of genes involved in bile acid transport and cholesterol homeostasis in the PD ileum (n=6 controls, 8 PD patients) and liver (n=6 controls, 6 PD patients). Ileum and liver samples from PD patients and healthy controls were matched for sex, age, and postmortem interval. Tissue (30-50mg per sample) was homogenized in 1 mL TRIzol with a handheld homogenizer for ileum and with Precellys bead tubes for liver. Following standard TRIzol RNA extraction, samples were treated with DNase (Qiagen) for 30 min. RNA cleanup was performed using a RNeasy column (Qiagen) according to the manufacturer’s instructions, with the addition of two 5 min washes with 75% ethanol. Isolated RNA quantity was determined with a NanoDrop 2000 spectrophotometer, and RNA integrity was confirmed with an Agilent 2100 Bioanalyzer. RNA was converted to cDNA using a High Capacity cDNA kit (Applied Biosystems). Samples were analyzed by qPCR with TaqMan reagents (Applied Biosystems; Supplementary File 11), using 25 ng of cDNA per qPCR reaction. Samples were run in triplicate and results were normalized to plate standardization controls. Delta delta CT values of gene transcripts were used to determine statistical changes in the ileum and liver of PD patients relative to controls, normalized to housekeeping control genes (β-actin and HPRT for liver, β-actin or villin for ileum). Statistical analysis was performed using one-way ANOVA with *p*-values <0.05 considered to be significant changes.

### Blood marker analysis in Parkinson’s Progression Markers Initiative (PPMI) data

We examined whether common blood markers of liver function were altered in PD patients and with the severity of the disease using the highly clinically characterized PPMI dataset (Parkinson Progression Marker Initiative, 2011). Data was downloaded from the PPMI website (https://www.ppmi-info.org/access-data-specimens/download-data/) on May 13 2020, and included screening demographics, blood chemistry hematology, PD features, MDS Unified Parkinson Disease Rating Scale (UPDRS) patient questionnaire, and prodromal diagnostic questionnaire. Five commonly used tests to check liver abnormalities, including alanine transaminase (ALT), aspartate aminotransferase (AST), alkaline phosphatase (ALP), albumin, and bilirubin, were examined. The MDS-UPDRS score was used to examine the relationship of blood markers with the severity of the disease. MDS-UPDRS total score was calculated based on PPMI score calculations standard by 59 variables from MDS-UPDRS part I, II and III, and PD motor symptoms were determined from MDS UPDRS part III. In our analysis, we examined the *de novo* PD cohort, which included 475 newly diagnosed PD patients (within 2 years of diagnosis) who were not taking PD medications at the time of enrollment and 231 healthy controls. PD patients had DaTscan imaging evidence of dopamine deficiency and at least two of the following: resting tremor, bradykinesia, and rigidity. We also analyzed the genetic PD cohort, which included 396 PD patients with a mutation in *LRRK2*, *GBA* or *SNCA* as well as 619 controls.

We first determined blood markers altered in *de novo* PD and in genetic PD patients, controlling for the age at the time of blood draw, sex, and anemia measures (levels of red blood cells, hemoglobin, and hematocrit as three additional covariates). In this analysis, averaged values for each blood marker of each individual in the PPMI was used, and statistical analysis was performed using a robust linear regression in the limma package (Ritchie et al., 2015). Next, we determined blood markers linked to the severity of the motor symptoms (as defined by MDS-UPDRS III) and total disease symptoms (MDS-UDPRS total score), adjusting for disease duration, age of PD onset, sex, and anemia measures, as well as UPDRS score and disease duration interaction. Statistical analysis was performed by a linear mixed-effects model in lme4 (Bates et al., 2015). Finally, we assessed differences in blood markers of liver function in the prodromal cohort. The prodromal cohort consisted of individuals with REM sleep behavior disorder (RBD) or hyposmia who are at risk of developing PD. Blood markers were evaluated at the date of enrollment into the PPMI, prior to any PD diagnosis. We separated the prodromal PD patients into the group of individuals that were later diagnosed with PD (n=36 converters) and those that were not (n=102 non-converters), comparing these groups with controls (n=561 controls). We also compare the blood markers of converters at the time of first PD diagnosis to those of controls, non-converters, and converters before PD diagnosis at the time of enrollment into the PPMI. Statistical analysis was performed using a robust linear regression in the limma package, adjusting for age at the time of blood draw, sex, and anemia measures. *P*<0.05 was the threshold for statistical significance.

## Supplementary Figures

**Figure S1.**
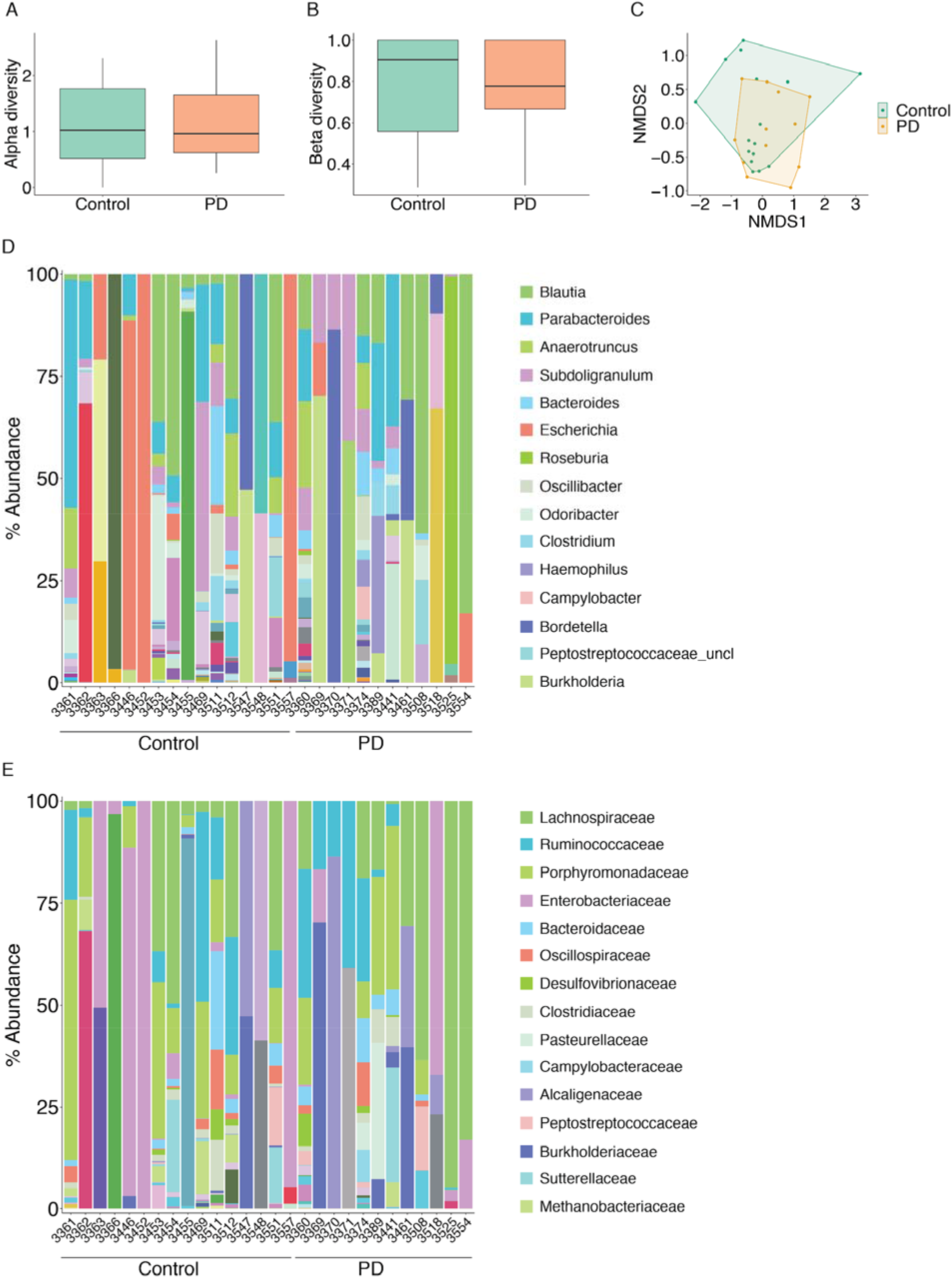
Microbial diversity in the human appendix. The functional microbiome was determined in metatranscriptomic sequencing data from 12 PD patients and 16 controls. (**A**) Alpha diversity (Shannon index) calculated by vegan package. (**B**) Beta diversity (Whittaker index) calculated by vegan package. (**C**) NMDS plot showing the distribution of samples according to the microbial community. (**D**) Microbiota composition in the human appendix at the genus level. Top 15 most abundant microbiota genera are listed. (**E**) Microbiota composition in the appendix at the family level. Top 15 most abundant microbiota families listed.

**Figure S2.**
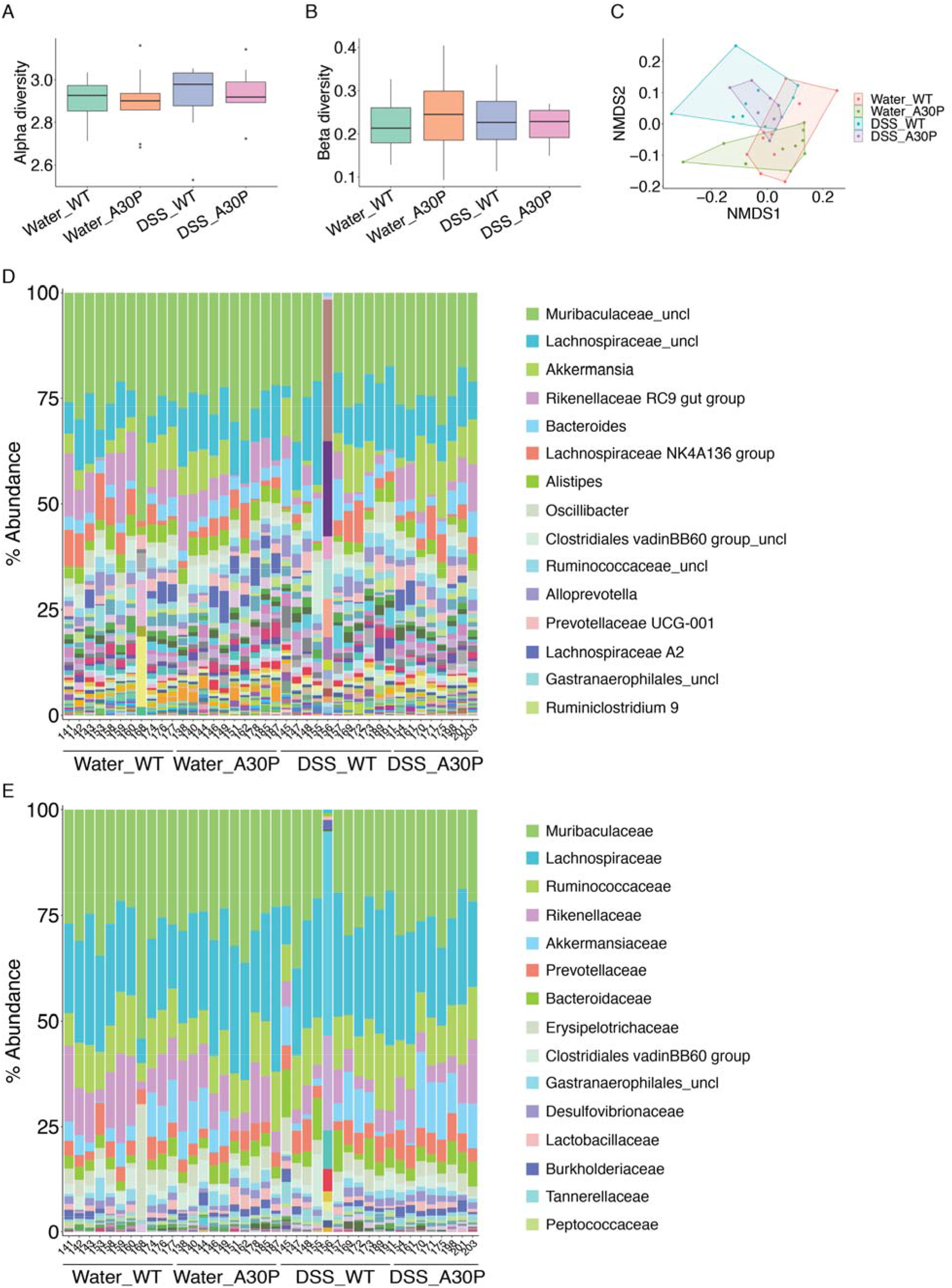
Microbial diversity in the mouse model of synucleinopathy and gut inflammation. The microbiome was profiled by 16S rRNA sequencing in the cecal patch of the A30P α overexpressing mice and wild-type mice previously exposed to DSS colitis or water. (**A**) Alpha diversity (Shannon index) calculated by vegan package. (**B**) Beta diversity (Whittaker index) calculated by vegan package. (**C**) NMDS plot showing the distribution of samples according to the microbial community. For visualization, outlying samples 168 and 156 are not shown in A, B, or C but were included in the analysis. (**D**) Microbiota composition in the mouse cecal patch at the genus level. Top 15 most abundant microbiota genera are listed. (**E**) Microbiota composition in the mouse cecal patch at the family level. Top 15 most abundant microbiota families listed.

**Figure S3.**
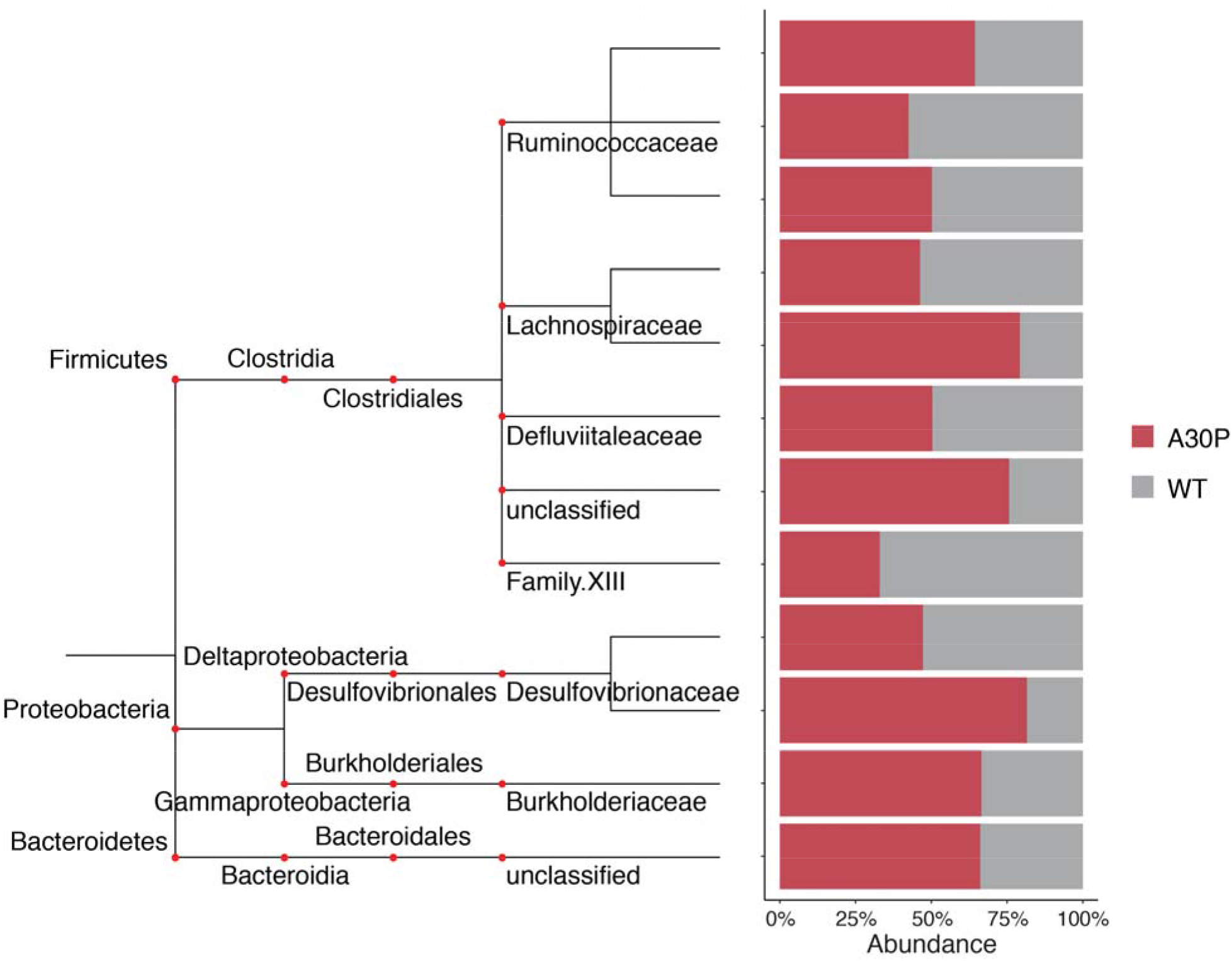
Microbiome alterations in the A30P. α**-syn mouse cecal patch.** Abundance (percentage) of each microbial genus altered in A30P α-syn overexpressing mice relative to wild-type mice with nominal *p*<0.05. Taxonomic tree denotes phylum to family level.

**Figure S4.**
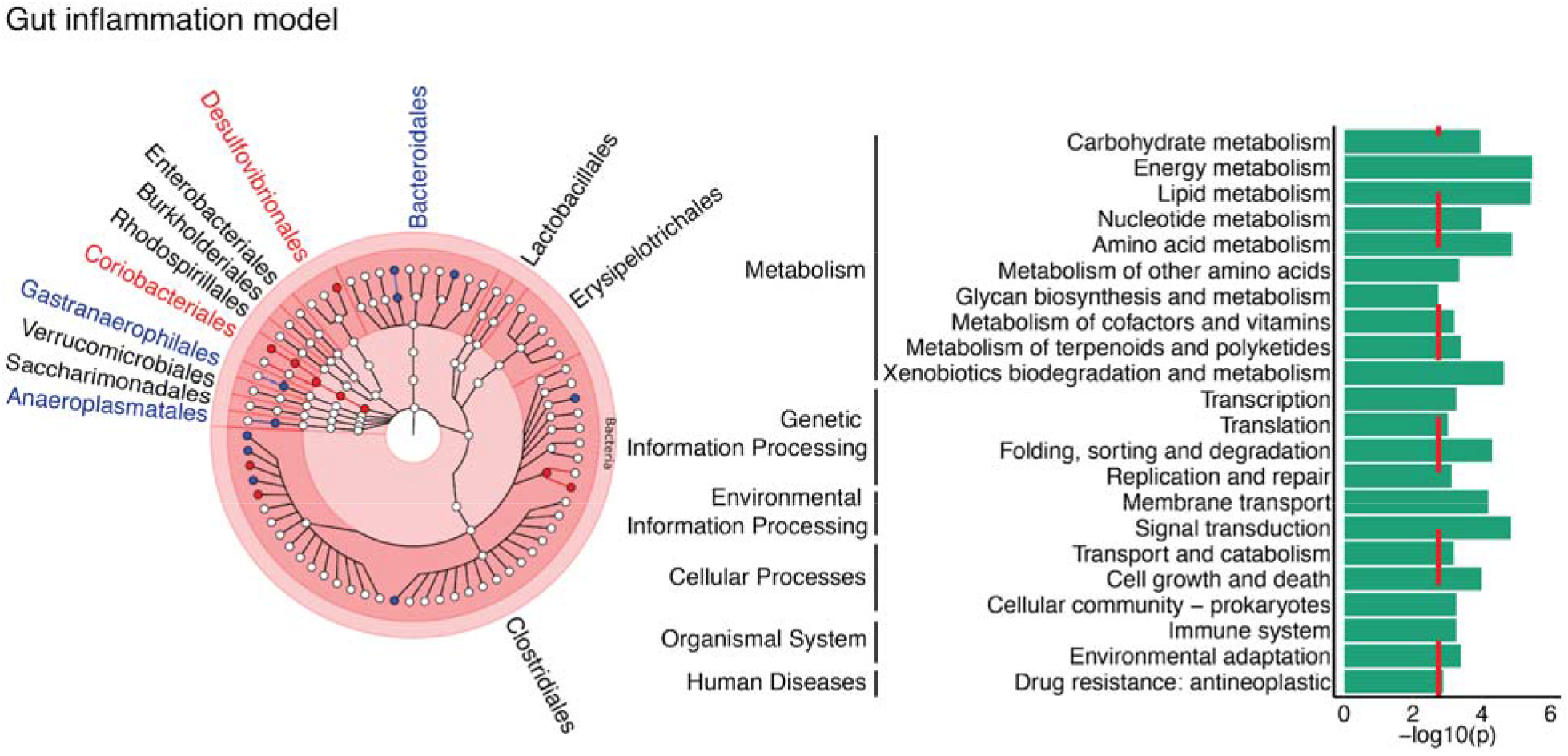
Microbiome changes in response to gut inflammation in the mouse cecal patch. Microbiome changes in mice previously exposed to DSS-mediated gut inflammation were determined by 16S rRNA sequencing. The 16S rRNA data were analyzed using QIIME2 and a zero-inflated gaussian mixture model in metagenomeSeq, adjusting for covariates (sex and genotype) was used to identify differences in the microbiome (*q*<0.05, metagenomeSeq). Microbial metabolic pathway changes were determined by PICRUSt2. n=21 water and 19 DSS colitis exposed wild-type and A30P α-syn mice. Taxonomic tree shows changes in microbial taxa (left) and related microbial functional pathways (right) in the DSS exposed mice, relative to mice given water. Taxonomic tree built using GraPhlAn, with kingdom in the center, and branching outwards to phylum, class, order, family, and genus. Microbial taxa highlighted in red are increased in the cecal patch in response to DSS colitis, relative to water, and blue are decreased. The red dashed line represents pathways that are *q*<0.1, by a zero-inflated gaussian mixture model in metagenomeSeq.

**Figure S5:**
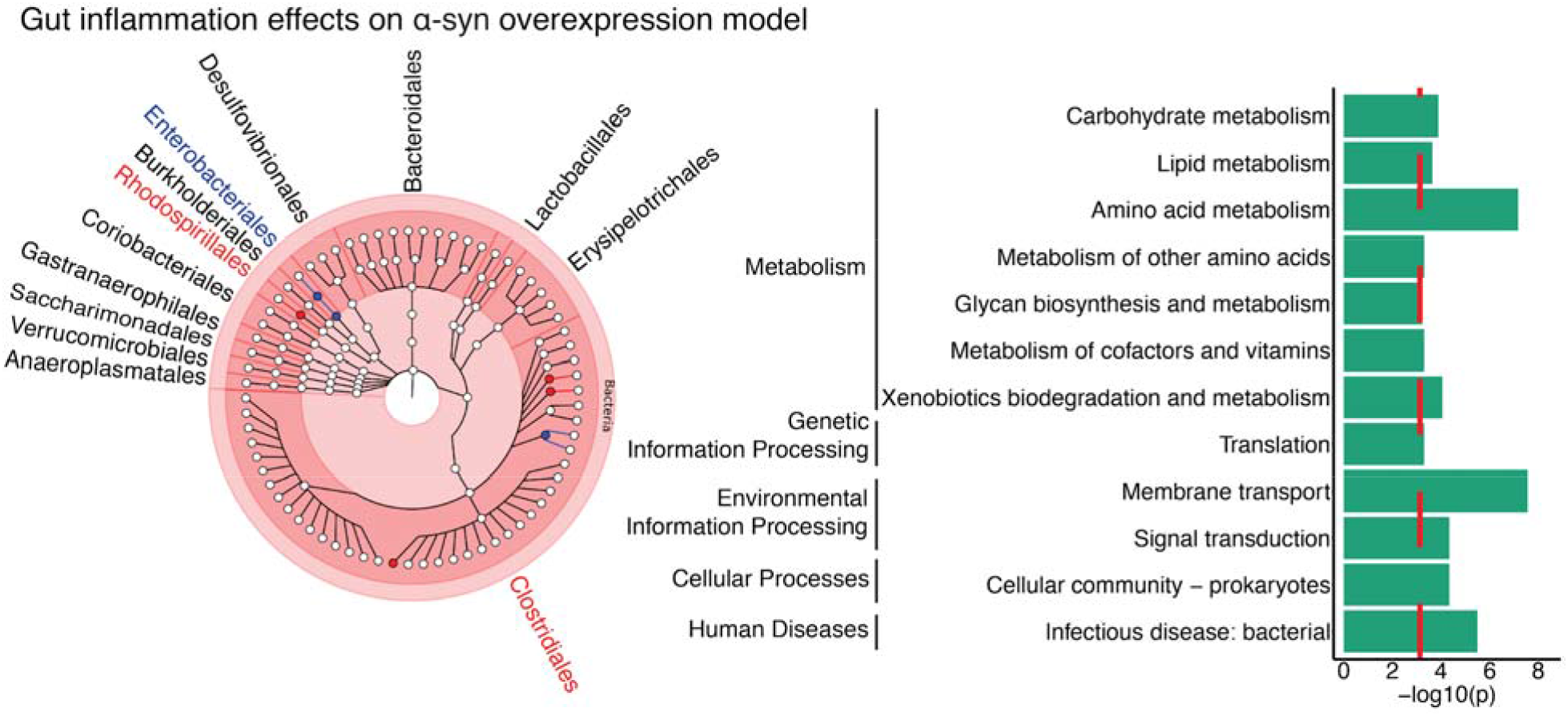
Microbial changes in the synucleinopathy model in response to gut inflammation. Microbiome (left) and related pathways (right) differentially altered in the cecal patch of A30P α-syn overexpressing mice in response to gut inflammation (n=8 A30P/DSS, 10A30P/water, 11 WT/DSS, 11 WT/water). The 16S rRNA sequencing data were analyzed with QIIME2 and pathway analysis was performed using PICRUSt2. Microbiome changes were determined by metagenomeSeq, followed by a contrasts.fit comparison in limma, adjusting for sex (*q*<0.05, metagenomeSeq). Taxonomic tree built using GraPhlAn, with kingdom in the center, and branching outwards to phylum, class, order, family, and genus. Microbial taxa highlighted in red are increased in the cecal patch of A30P α-syn mice in response to DSS colitis, and blue are decreased. Microbial functional pathways differentially altered in A30P α-syn mice following DSS colitis are shown, and red dashed line represents *q*<0.1 pathways.

**Figure S6:**
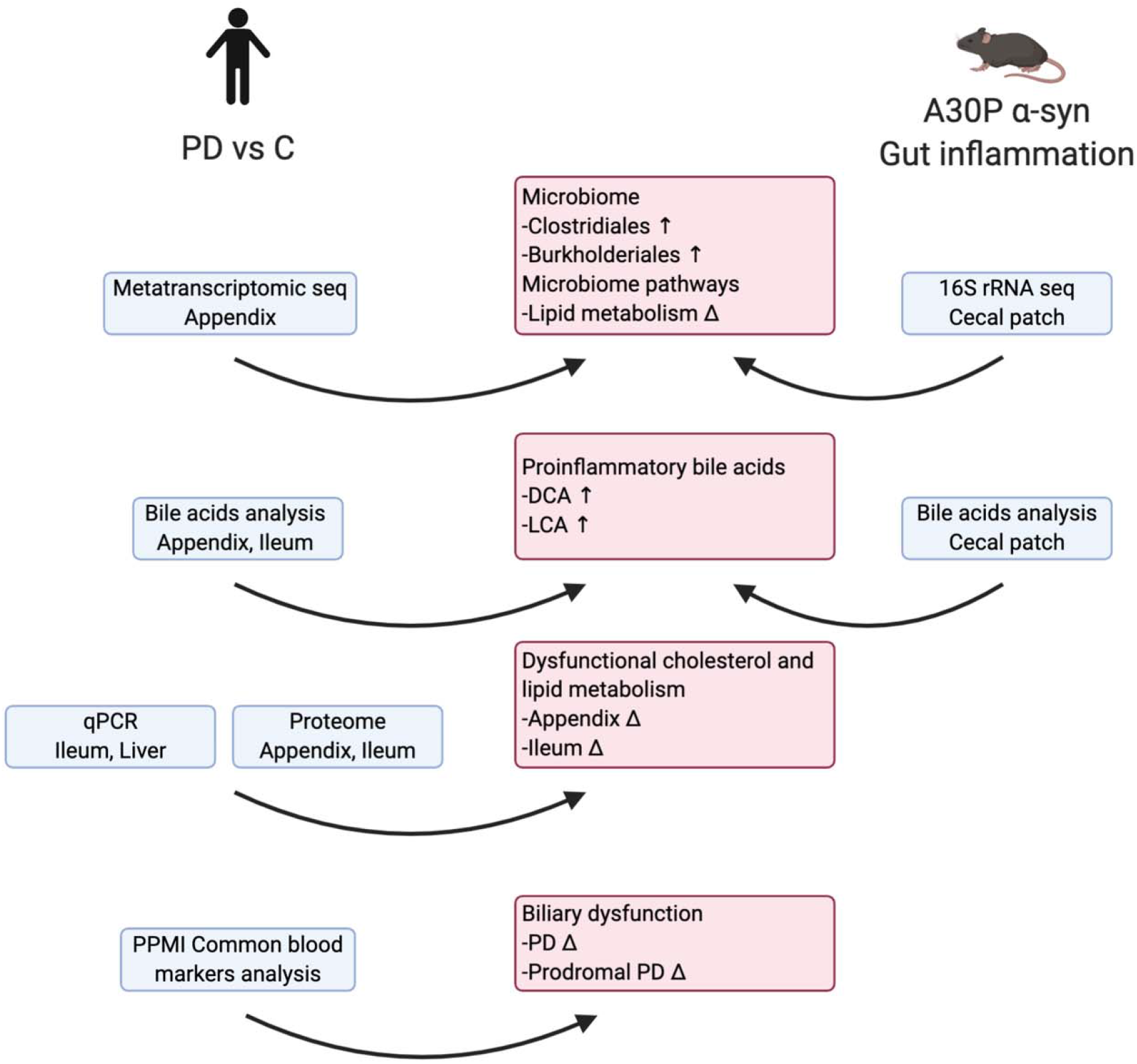
Experimental design and main findings of our study investigating functional changes in the appendix microbiome in PD. The human appendix is an immunological organ that is also considered to be a storehouse for the gut microbiome. Recently, the appendix has been implicated in the risk of developing PD (Killinger et al., 2018). To determine whether the PD appendix exhibits functional changes in the microbiome, we performed a metatranscriptomic analysis of the PD and control appendix. Microbiome changes in PD were compared to those in the cecal patch (appendix-equivalent) of a mouse model of synucleinopathy and gut inflammation. We identified microbiome changes in the PD appendix that affect lipid homeostasis and the synthesis of secondary bile acids, which in turn led us to an analysis of bile acids. In the PD appendix, we found an increase in the microbially-derived, cytotoxic bile acids LCA and DCA. These bile acid changes were recapitulated in the mouse model of synucleinopathy. Proteomic and transcript analysis also demonstrated a disruption in cholesterol and lipid metabolism in the PD gut. Since elevated LCA and DCA can lead to liver injury and gallstone disease, we profiled blood markers of liver and gallstone dysfunction in PD patients using the PPMI dataset. We found evidence of biliary dysfunction in PD patients, which occurred even before clinical onset of motor symptoms.

## Supplementary Files

Supplementary File 1. Demographic and clinical information for human samples.

Supplementary File 2. Differences in PD appendix microbiome and related pathways identified by metatranscriptomic analysis

Supplementary File 3. Mouse sample information.

Supplementary File 4. Differences in A30P α-syn mouse cecal patch microbiome and related Pathways

Supplementary File 5. Differences induced by gut inflammation in the mouse cecal patch microbiome and related pathways

Supplementary File 6. Combined effect of synucleinopathy and DSS inflammation on the mouse cecal patch microbiome and related pathways

Supplementary File 7. Proteomic changes in the human PD appendix and ileum.

Supplementary File 8. Changes in bile acid composition in the human PD appendix, and ileum and in the cecal patch of mice with synucleinopathy and gut inflammation

Supplementary File 9. Differences in gene transcripts related to bile acid homeostasis in the human PD ileum and liver identified by qPCR

Supplementary File 10. Changes in blood markers of biliary function in PD patients in the PPMI data

Supplementary File 11. Primers used in liver and ileum qPCR experiment

